# The impact of chromatin remodeling on gene expression at the single cell level in *Arabidopsis thaliana*

**DOI:** 10.1101/2020.07.27.223156

**Authors:** Andrew Farmer, Sandra Thibivilliers, Kook Hui Ryu, John Schiefelbein, Marc Libault

**Affiliations:** National Center for Genome Resources, Santa Fe, New Mexico, USA; Department of Agronomy and Horticulture, Center for Plant Science Innovation, University of Nebraska-Lincoln, Beadle Center, Lincoln, Nebraska, USA; Department of Molecular, Cellular, and Developmental Biology, University of Michigan, Ann Arbor, MI, USA

## Abstract

Similar to other complex organisms, plants consist of diverse and highly specialized cell types. The gain of unique biological functions of these different cell types is the consequence of the establishment of cell-type-specific transcriptional programs and their associated regulatory mechanisms. Recently, single cell transcriptomic approaches have been applied on *Arabidopsis thaliana* root protoplasts allowing the accurate characterization of the transcriptional profiles of the cell-types composing seedling roots. As a first step in gaining a deeper understanding of the regulatory mechanisms controlling Arabidopsis gene expression, we report the use of single nucleus RNA sequencing (sNucRNA-seq) and single nucleus Assay for Transposase Accessible Chromatin sequencing (sNucATAC-seq) technologies on Arabidopsis roots. The comparison of our single nuclei transcriptomes to previously published protoplast transcriptomes validated the use of nuclei as biological entities to establish cell-type specific transcriptomes from multicellular organs. Furthermore, our sNucRNA-seq results uncovered the transcriptome of additional cell subtypes not identified by scRNA-seq. Similar to our transcriptomic approach, the sNucATAC-seq approach led to the distribution of the Arabidopsis nuclei into distinct clusters suggesting the differential remodeling of the chromatin between groups of cells according to their identity. To reveal the impact of chromatin remodeling on gene transcription, we integrated sNucRNA-seq and sNucATAC-seq data and demonstrated that cell-type-specific marker genes also display cell-type-specific pattern of chromatin accessibility. Our data suggest that the differential remodeling of the chromatin is a critical mechanism to regulate gene activity at the cell-type level.

## Introduction

The gain of unique biological functions by the various cell-types composing a plant depends on their differential use of the same genomic information to produce cell-type-specific transcriptional profiles. The differential use of the genomic information between cells and cell-types is thought to rely, in part, on the differential remodeling of the chromatin. Changes in the landscape of chromatin fiber influences the accessibility of the genomic DNA for regulatory proteins such as transcription factors. This latter statement is supported by the human ENCODE project, which recently revealed that the establishment of a chromatin landscape at the single cell level was highly informative to reveal putative TF-binding sites [1]. Ultimately, each plant cell will activate or repress specific sets of genes to fulfill the biological functions inherent to their cell-type and their response to environmental stresses. In animal science, single cell RNA-seq (scRNA-seq) and ATAC-seq [2] technologies have been successfully applied across various cell-types and tissues to better understand the impact of the dynamic remodeling of the chromatin on gene expression [3–6]

Recently, scRNA-seq approaches have been applied to Arabidopsis root protoplasts, allowing the accurate characterization of the transcriptional profiles of thousands of cells and their differential regulation in mutants or in response to a stress [7–11]. These studies revealed the power of single cell technologies to establish the transcriptomic maps of various Arabidopsis root cells and cell-types and the dynamic regulation of gene expression during cell development. However, using plant protoplasts as biological entities to analyze gene expression is problematic. For example, some cell types are resistant to cell wall digestion [e.g., stele cells [7–11]], the protoplasting procedure itself has a significant impact on gene expression [7], and there is a bias toward sequencing of smaller-sized cells/protoplasts [7]. In addition, the effective isolation of plant protoplasts requires the development of particular cell-wall degrading enzymatic cocktails tailored for the differential biochemical composition of the cell wall existing between plant species, the developmental stage of the cell (i.e., differential biochemical composition between the primary and secondary cell wall), and the relative position of the root cell (i.e., external *vs.* internal locations) [12, 13].

As an alternative to protoplasts, bulks of plant nuclei have been used to gain transcriptomic information from plant cells. For instance, transcriptomes were established from nuclei populations using Isolation of Nuclei from TAgged specific Cell Types (INTACT) technology on rice roots [14], Arabidopsis embryeso [15], and seed endosperm [16]. Also, taking advantage of the nuclear ploidy of the cells composing the pericarp tissue of the developing tomato fruits, Pirello et al., (2018) analyzed the expression of tomato genes using a population of nuclei characterized by various levels of endoreduplication [17]. Nevertheless, these methods also suffer from various limitations. For instance, the use of the INTACT technology presupposes the identification of cell-type-specific marker genes to express a reporter gene and require the generation of transgenic material.

To overcome some of the challenges associated with the generation and manipulation of plant protoplasts, we tested the use of isolated plant nuclei to establish the transcriptome of thousands of plant single cells. Previous single nuclei transcriptomic experiments conducted on various animal systems suggest that nuclei could be used to establish biologically meaningful transcriptomic information compared to isolated cells [18, 19]. Our data reveal high similarities existing between the nuclear and protoplast transcriptomes and the discovery of additional root cell-type transcriptomes that support the use of isolated nuclei as valuable biological entities to access single cell gene expression. To gain a deeper understanding of the regulatory mechanisms controlling gene expression in and between Arabidopsis root cells and cell-types, we also developed and applied single nuclei ATAC-seq (sNucATAC-seq). SNucATAC-seq revealed cell-type-specific remodeling of the chromatin and dynamic changes occurring in chromatin remodeling during cell differentiation and development. Upon establishing the unique and conserved chromatin landscapes for each major Arabidopsis root cell type, we integrated sNucRNA-seq and sNucATAC-seq datasets to identify promoter/enhancer sequences that likely regulate expression of nearby genes in a cell-type-specific manner, depending on the accessibility of the chromatin fiber. Altogether, we propose a set of “rules” governing the influence of chromatin accessibility on Arabidopsis gene activity.

## Results and Discussion

### Arabidopsis root nuclei generate biologically meaningful transcriptomic datasets

Nuclei from Arabidopsis seedling roots were purified and used with the 10X Genomics Chromium platform to create sNucRNA-seq libraries (Supplemental Figure 1). To accurately compare sNucRNA-seq and scRNA-seq transcriptomes, the Arabidopsis seedlings were grown and primary roots were isolated as described by Ryu et al., (2019) [9]. Across five independent biological replicates, we sequenced the transcriptome of 10,608 nuclei (See Supplemental Table 1 for sequencing details). Because some nuclear transcripts might not be spliced, we applied a “pre-mRNA” strategy to include intron-mapping reads. We obtained a mean of 1,126 expressed genes per nucleus, allowing the identification of 24,740 expressed genes out of the 27,416 predicted Arabidopsis protein-coding genes (90.2%). As a comparison, the transcriptome of 7,473 selected Arabidopsis protoplasts by Ryu et al., (2019) [9] allowed the detection of 4,751 expressed genes per cell and a total of 25,371 expressed genes (92.5%). We suggest that the higher number of expressed genes identified per protoplasts *vs.* nucleus is the consequence of the larger and more complex pool of polyadenylated transcripts in one cell *vs.* one nucleus, due to the accumulation of transcripts and their relative stability (i.e., the half-life of the cellular mRNA is estimated at 9 hours in human cells [20]). This conjecture also suggests that the nuclear transcriptome represents a snapshot of the dynamic transcriptional activity of the genes while the cellular transcriptome may represent an integration of gene activity overtime.

To evaluate the biological significance of the nuclear transcriptomes obtained from the sNucRNA-seq data, we performed correlation analyses between bulk RNA-seq from intact whole roots and from protoplast suspensions, and pseudo-bulk RNA-seq from root protoplasts and nuclei that were processed through scRNA-seq [9] and sNucRNA-seq technologies (present study). The transcriptome of intact whole roots is highly correlated with the transcriptome of a suspension of protoplasts and with both pseudo-bulked scRNA-seq and sNucRNA-seq datasets (i.e., Spearman’s Rank Correlation Coefficient = 0.902, 0.886, and 0.876 for all genes, = 0.905, 0.889, 0.878 upon excluding known protoplast-responsive genes [21], respectively). These results reveal that the sNucRNA-seq transcriptome of the Arabidopsis root is as highly correlated to a whole root transcriptome than protoplast-based transcriptomes. Overall, single nuclei-based transcriptome reflect well the transcriptome of a complex organ such as the Arabidopsis root.

Taking advantage of the capability of the Seurat package to integrate independent datasets [22, 23], we co-clustered 10,608 Arabidopsis root nuclei with 7,473 Arabidopsis root protoplasts [9] according to their transcriptomic profiles. Applying Uniform Manifold Approximation and Projection (UMAP), a dimensionality reduction technique which enhanced the organization of nuclei/cell clusters compared to tSNE [24], the Arabidopsis root nuclei and protoplasts were clustered into 21 and 18 different groups, respectively (Figure 1A). UMAP visualization clearly reveals the overlapping distribution existing between the 18 co-characterized clusters and confirms the identification of three new clusters upon applying sNucRNA-seq technology (clusters #16 and 19 include over 97% of nuclei and only 3% of protoplasts; while cluster #20 is exclusively composed by nuclei; Figure 1A, red circles).

**Figure 1.**
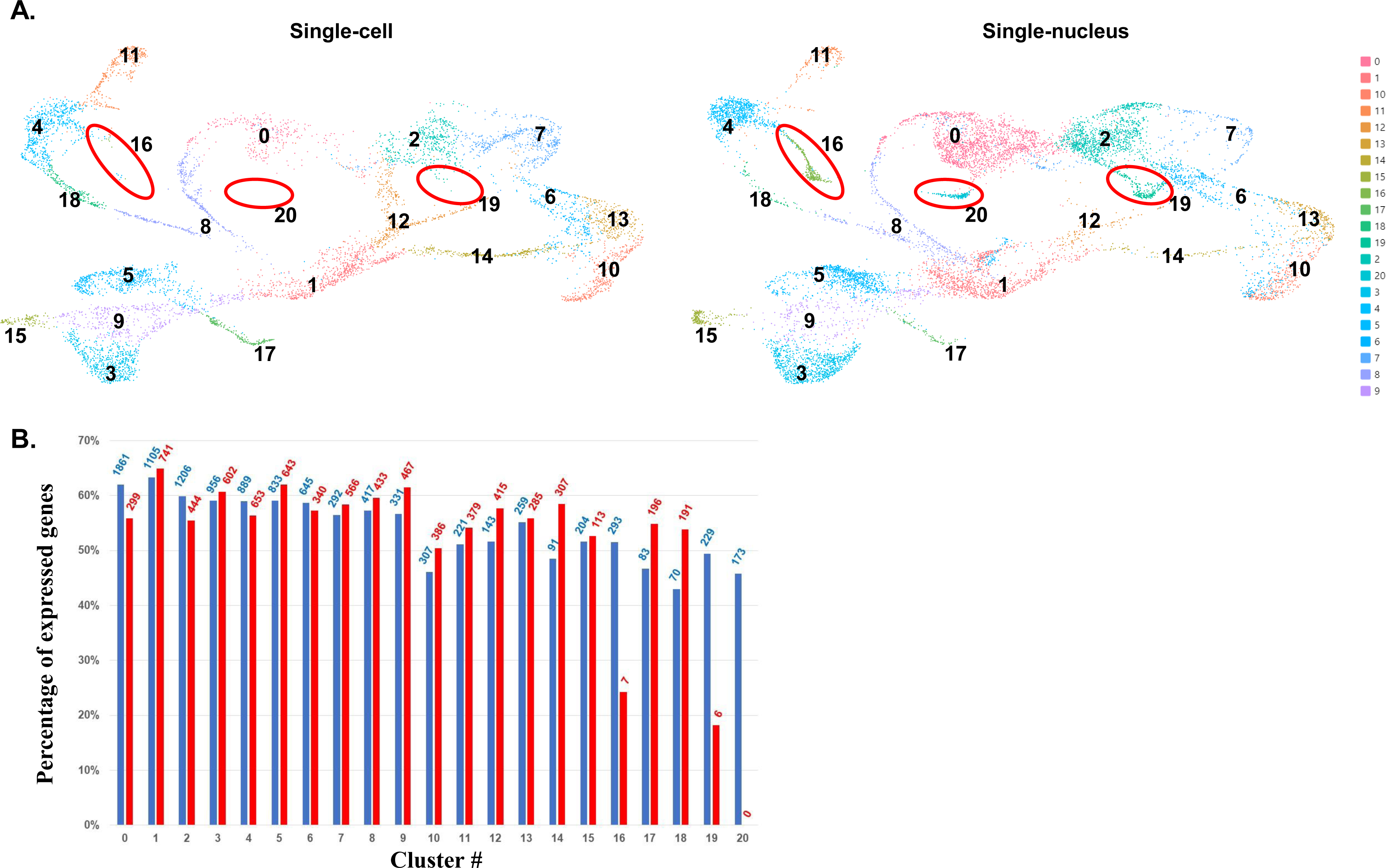
UMAP Cluster analysis of the Arabidopsis single-cell and single-nucleus transcriptomes. **A.** Comparative analysis of the clustering of Arabidopsis cells (left, [9]) and nuclei (right, present study) according to their transcriptomic profiles. Red circles highlight the clusters that are mostly or exclusively revealed using sNucRNA-seq technology. **B.** Percentage of Arabidopsis genes found expressed upon analysis of sNucRNA-seq (blue) and scRNA-seq (red) technologies for each 21 clusters. The numbers at the top of the bars highlight the number of nuclei and cells associated to each cluster.

To further evaluate the biological relevance of the sNucRNA-seq approach, we compared the percentages of protein-coding genes found expressed or not expressed using sNucRNA-seq and scRNA-seq technologies across the 18 co-characterized clusters (Figure 1B). Among the 27,416 Arabidopsis protein-coding genes, 30.9 (cluster #1) to 44.8% (cluster#10) were found not to be expressed by both sNucRNA-seq and scRNA-seq technology while 40.6% (cluster #18) to 59.1% (cluster #1) were found expressed by applying both technologies (Supplemental Figure 2). Only a small percentage of genes were found expressed using only one technology or the other [i.e., an average of 4.9% and 7.4% of the Arabidopsis genes were found expressed using sNucRNA-seq or scRNA-seq, respectively (Supplemental Figure 2)]. Taken together, these results revealed that over 80% of the expressed Arabidopsis protein-coding genes were detected by both sNucRNA-seq and scRNA-seq (Supplemental Figure 3). Excluding the clusters that are unique to sNucRNA-seq, the percentage of expressed genes per cluster identified using sNucRNA-seq technology (from 42.9 to 63.3 %) is not significantly different compared to the percentage of expressed genes per cluster identified using scRNA-seq technology (from 50.5 to 64.9%) (i.e., Student t test: P>0.136; Figure 1B). These results show that the cellular and nuclear transcriptomes provide similar transcriptomic information and suggest that isolated plant nuclei can be utilized to establish meaningful transcriptomic information at the single cell-type level. Through the identification of three new cell clusters, our data also suggest that the sNucRNA-seq approach captures a more diverse and representative population of Arabidopsis root cell-types compared to scRNA-seq.

### Functional assignment of Arabidopsis root cell clusters

Taking advantage of recently published Arabidopsis root single cell transcriptomes [7–11] and the current bibliography [25–28], we created a list of 103 cell-type marker genes (Supplemental Table 2). To assign biological entities to the 21 Arabidopsis clusters, we looked for the accumulation of transcripts for these marker genes (Figure 2A and B; Supplemental Figure 4). This strategy allowed us to characterize five major groups of cells: trichoblasts (clusters #6, 10, 13, and 14), atrichoblasts (clusters #1, 2, 7, and 12), cortex (clusters #0, 8, 16, and 20), endodermal (clusters #4, 8, 11, 16, and 18), and stele cells (clusters #3, 5, 9, 15, and 17) (Figure 2A). In addition, we observed the accumulation of transcripts of marker genes of the quiescent center in the cells and nuclei composing the cluster #1. We were also able to more specifically assign the different cell types composing the stele and characterize the developing *vs.* mature trichoblasts and endodermal cells (Figure 2A).

**Figure 2.**
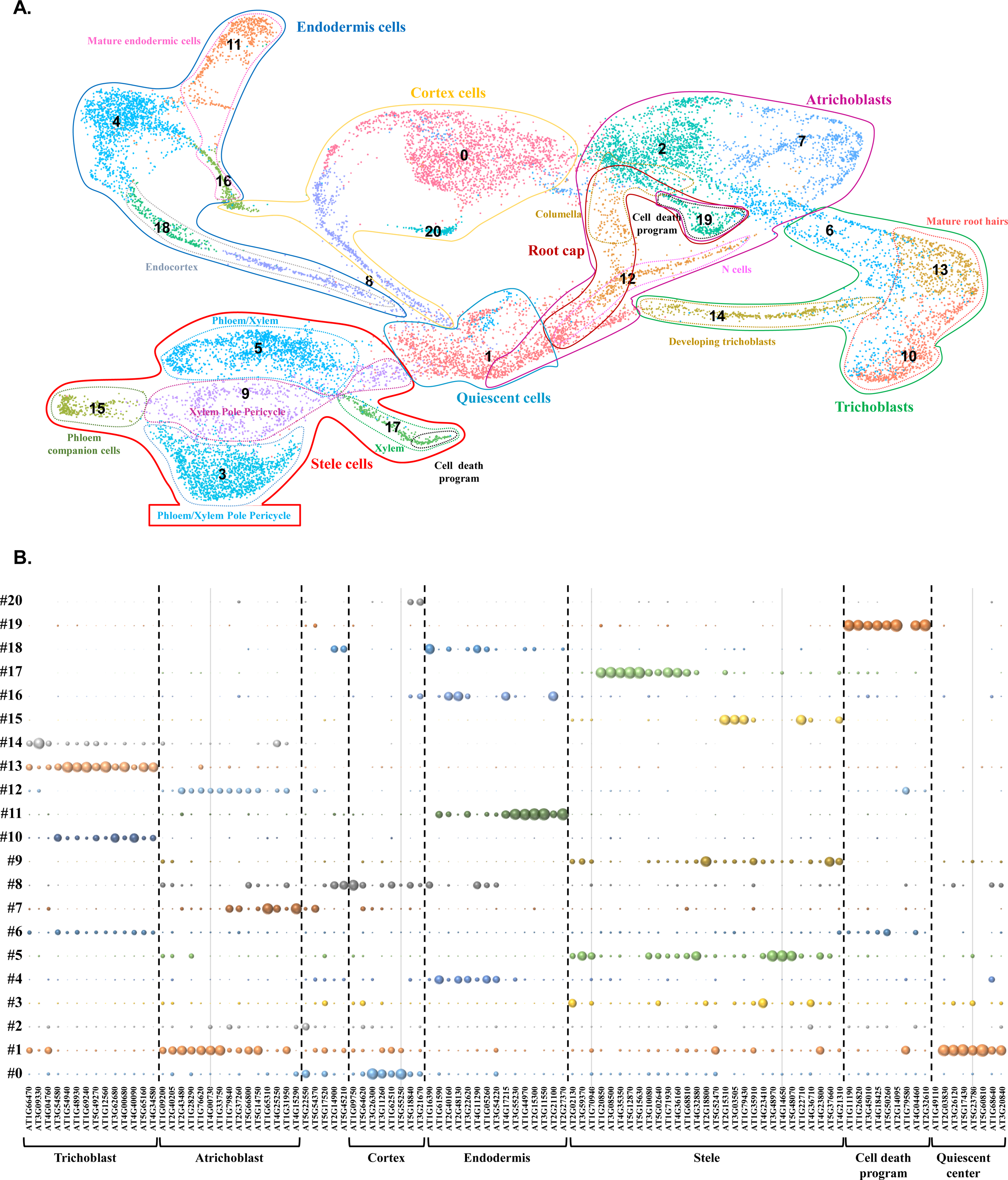
A. Assignment of Arabidopsis root cell-types based on the characterization of the expression profile of cell-type and cell death marker genes. **B.** Normalized expression level of 103 cell-type and cell death marker genes (x-axis, see Supplemental Table 2) across the 21 different clusters (y-axis). sNuc-RNAseq dataset were used to create this figure. The diameter of each circle reflects the relative expression of the genes in each cell cluster. As a comparison, the transcriptional activity of these genes in Arabidopsis protoplasts [9] is provided in Supplemental Figure 4.

The topology of the clusters generated by the UMAP technique reveal the functional organization of the cells and nuclei in and between the cell-types compared to previous reports using tSNE to cluster Arabidopsis protoplasts [8, 11]. For instance, the undifferentiated cells of the roots found in cluster #1 are localized in the center of the UMAP map. Originated from cluster #1, several elongated projections of cells (e.g., clusters #8, 12, 14, and 18) end with more globular clusters (e.g., clusters #0, 2. 3, 4, 5, 7, 9, 10, and 13). The elongated clusters likely reflect the progressive changes occurring in the transcriptomic programs during cell differentiation whereas the globular clusters represent the differentiated cells composing the Arabidopsis roots.

Among the three clusters specifically defined using sNucRNA-seq technology (i.e., clusters #16, 19 and 20; Figures 1A and 2A), cluster #16 could be divided into two different sub-groups which are characterized by the activity of endodermal and cortex marker genes, respectively [i.e., the AT1G61590 (*PBL15*), AT2G40160 (*TBL30*), AT2G48130, and AT4G17215 as endodermal marker genes; the AT5G18840 and AT3G21670 (*NPF6.4/NRT1.3*) as cortical marker genes; Supplemental Figure 5 (red and blue circles, respectively); Supplemental Table 2]. This observation is also supported by the relative UMAP localization of the cluster#16 that connects the endodermis and cortex cell clusters (Figure 2A). Cluster #16 is also characterized by the specific expression of genes encoding peroxidases (e.g., AT1G68850, AT2G35380) and GDSL-motif esterase/acyltransferase/lipase (e.g., AT2G23540, and AT5G37690 [29]) (Supplemental Figure 5, blue arrows). Members of the *GDSL* family such as the rice *WDL1* and the tomato *GDSL1* genes control the process of cellular differentiation [30–32] suggesting that the cluster #16 might contain differentiating cells. This hypothesis is supported by the role of root peroxydases in controlling the production of reactive oxygen species to regulate cell elongation and differentiation [33]. Interestingly, the *UPBEAT1* gene (AT2G47270) has been identified as a major repressor of the transcriptional activity of peroxidases genes and ROS distribution, and a negative regulator of the size of the Arabidopsis root apical meristem by modulating the balance between cell proliferation and differentiation [33]. Mining the UMAP clusters, we found that *UPBEAT1* is broadly expressed at the exception of the clusters #10, 16, 19, and 20 (Supplemental Figure 5, black arrows). GDSL lipases are also playing a central role in cutin biosynthesis [34]. Mining the sc/sNucRNA-seq datasets, we identified many other genes preferentially expressed in cluster #16 and involved in the biosynthesis of suberin and cutin [e.g. α/β hydrolases (AT4G24140), *GPAT5* that plays a central role in suberin biosynthesis (AT3G11430) [35, 36], AT1G49430, AT2G38110, *GPTA4* (AT1G01610) and *8* (AT4G00400) [37], and another GDSL-like lipase (AT1G74460) [37]; Supplemental Figure 5 (blue arrows)]. Previous studies revealed that suberin and cutin are notably deposited at the location of the emergence of lateral roots [38]. Taken together, the transcriptional pattern of the *UPBEAT1*, peroxidases, *GDSL* genes, and other suberin/cutin biosynthesis-related genes, associated with the activity of both cortical and endodermal marker genes suggest that the cells associated with the cluster #16 are associated with the emerging lateral roots.

Cluster #19 (Figure 2A) is characterized by the specific expression of *CEP1* (AT5G50260) and *EXI1* (AT2G14095) (Supplemental Figure 6), two genes previously characterized as regulators of the cell death program in the root cap [27]. In addition, we identified *KIRA1* (AT4g28530), a gene controlling cell death during flower development [39], as specifically expressed in cluster #19. Additional cell death marker genes (i.e., *BFN1*, *RNS3*, *SCPL48*, *DMP4*, and *PASPA3*) were also found specifically expressed in cluster #19 (Figure 2B) and in a subset of the xylem cluster #17 [27]. Previous studies showing that the cell death programs play critical roles in the development of the xylem and root cap [40, 41] support the specific activity of these cell death marker genes in the xylem cluster #17 and the assignment of the cells composing cluster #19 as root cap cells. Interestingly, we also noticed the specific expression of two genes encoding α/β hydrolases in cluster #19 (e.g., AT4G18550, AT1G73750; Supplemental Figure 6). We hypothesize that these genes might play a role in the accumulation of cutin and suberin in the external layer of the Arabidopsis root cap [38]. The characterization of the transcriptome of the cells in the clusters#16 and 19 from isolated nuclei and not from isolated protoplasts could be due to the low digestibility of the cell wall of these cells, a consequence of the accumulation of suberin and cutin.

Cluster #20 (Figure 2A) is characterized by the expression of a subset of the cortex-specific marker genes (e.g., AT5G18840 and AT3G21670 (*NPF6.4/NRT1.3*); Supplemental Figure 7; Supplemental Table 2; [7–9]). Among the genes specifically expressed in this cluster, *SCRAMBLED/STRUBBELIG* (*SCM*, AT1G11130) plays a critical role in the patterning of the root epidermal cells [42] (Supplemental Figure 7). The specific clustering of the *SCRAMBLED/STRUBBELIG*-expressing cells is further supported by the expression of several genes involved in lipid metabolism in cluster #20 [e.g., AT1G45201 (*TLL1*; triacylglycerol lipase-like 1); AT5G63560 (HXXXD-type acyltransferase); Supplemental Figure 7]. Membrane lipid remodeling has been shown to also play a critical role in root hair cell differentiation [43, 44]. Taken together, the cortical cells composing cluster #20 are hypothesized to play a role in the differentiation and patterning of the Arabidopsis epidermal root cells.

### Single-cell resolution ATAC-seq reveals the impact of chromatin folding on gene expression

While genomic information is almost identical between somatic cells (i.e., with the exception of somatic mutations), its differential use, notably through differential chromatin accessibility between cells, is required in order to fulfill their unique biological function through cell-type specific transcriptional regulation of the genes [45–48]. To date, bulk RNA- and ATAC-seq datasets have shown low correlations [45]. This could be the consequence of the cellular heterogeneity of the samples used. This hypothesis is supported by the human ENCODE project, which recently revealed that the establishment of a chromatin landscape at the single cell level was highly informative to reveal putative TF-binding sites [1]. To better evaluate the impact of chromatin remodeling in controlling plant gene expression between cells and cell types, we applied 10x Genomics sNucATAC-seq technology on 6768 Arabidopsis root nuclei isolated from two independent biological replicates.

Upon paired-end sequencing, a median of 10,253 independent genomic DNA fragments per nucleus were mapped against the Arabidopsis genome and a total of 20,803 accessible sites were characterized. As a comparison, Lu et al., (2017) and Tannenbaum et al., (2018) identified around 20,000 and 40,000 accessible sites from bulk ATAC-seq analyses conducted on Arabidopsis seedling and roots, respectively [49, 50]. As previously reported [49], we observed that the accessible regions of the chromatin are mostly located near transcription start sites (TSSs) where are located cis-regulatory elements (Figure 3A). Hypothesizing that each Arabidopsis root cell type is characterized by discrete and unique profiles of the remodeling of the chromatin fiber to control the activity of cell-type-specific genes, we applied the UMAP algorithm to cluster the 6768 nuclei into 21 different groups according to their chromatin folding profiles (Figure 3B).

**Figure 3.**
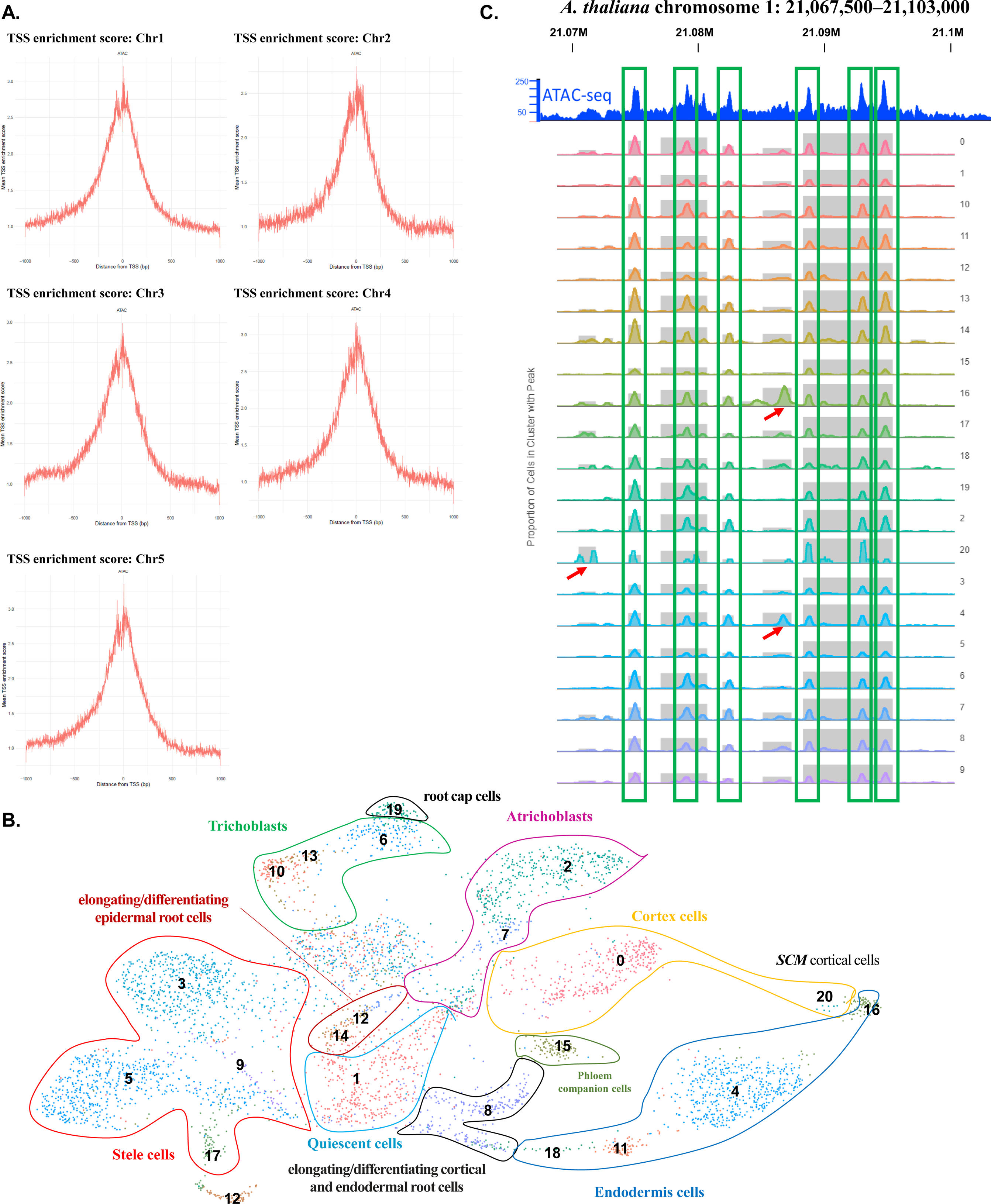
Use of sNucATAC-seq technology to characterize the differential folding of the chromatin fiber between Arabidopsis root cell types. **A.** Distribution of accessible regions of the chromatin fiber located around the TSS of the genes on the 5 Arabidopsis chromosomes after applying sNucATAC-seq technology. **B.** Clustering of Arabidopsis root nuclei upon sNucATAC-seq analysis. The numerical annotation of these clusters is the same as for the sNucRNA-seq clusters (Figure 2A). Taking advantage of the integrative sNucATAC-seq and sc/sNucRNA-seq analysis, we functionally assigned each of the 21 sNucATAC-seq Arabidopsis root clusters. **C.** Comparative analysis of the distribution of ATAC-seq peaks on the *A. thaliana* chromosome 1: 21,067,500–21,103,000. The top blue graphic was generated from a bulk ATAC-seq analysis of the Arabidopsis root seedling [55]. Below is indicated the ATAC-seq peak profiles across the 21 sNucATAC-seq clusters. The red arrows highlight cluster-specific sNucATAC-seq peaks.

To better estimate the gain in resolution of sNucATAC-seq versus bulk ATAC-seq datasets and to estimate the potential of sNucATAC-seq technology to reveal discrete changes in chromatin remodeling, we first compared the sNucATAC-seq and bulk ATAC-seq profiles generated from Arabidopsis root nuclei at one locus (i.e., chr1: 21,067,500–21,103,000) [50]. Across the 21 clusters, we can clearly identify the same major peaks revealed by bulk ATAC-seq technology [50] (Figure 3C, green boxes). Furthermore, our sNucATAC-seq approach revealed additional peaks but only in a subset of the 21 clusters (Figure 3C, red arrows). This first targeted analysis suggests that a single cell resolution ATAC-seq analysis has the potential to reveal discreet and cell-type-specific loci of accessible chromatin. These results also support the idea that the 21 sNucATAC-seq clusters are characterized by unique combinations of open chromatin loci. We conducted further analyses to reveal how cell-type chromatin accessibility play a critical role in controlling gene expression.

Considering that chromatin accessible sites are pre-required to promote gene expression, we assume that cell-type-specific ATAC-seq peaks located nearby the TSS contribute to the regulation of expression of cell-type marker genes (i.e., the peaks have at least one base pair overlap with the annotated transcripts, and extend upstream). Accordingly, we searched for correlations between the presence/absence of open chromatin nearby the TSS of genes and their transcriptional activity by integrating sNucATAC-seq and sc/sNucRNA-seq analysis using the Signac package v0.2.5 [51]. This strategy led to the annotation of the 21 sNucATAC-seq clusters similarly to the sc/sNucRNA-seq clusters (i.e., from 0 to 20; Figure 3B). We noticed that the sNucATAC-seq root cell clusters share a similar topography as the 21 sNuc/scRNA-seq clusters. First, the sNucATAC-seq UMAP clusters #6, 10, and 13; cluster #7; clusters #3, 5, 9, and 17; clusters #4, 11, 16, and 18; and clusters #0 and 20 which are composing the trichoblasts, the atrichoblasts, and the stele, endodermal, and cortical cells, respectively, are grouped in super-clusters which are similar to the sNucRNA-seq super-clusters (Figures 2A and 3B). Second, we noticed that, similar to the organization of the sc/sNucRNA-seq clusters, the sNuc-ATAC-seq clusters associated with undifferentiated and differentiating cells (i.e., clusters #1, 8, 12 and 14) are located in the center of the sNucATAC-seq UMAP map while the clusters associated with differentiated cells are located at its periphery. These similar topographies suggest that chromatin remodeling and gene expression could be both used as molecular markers to annotate plant cell-types and support the correlation existing between chromatin folding and the transcriptional activity of the Arabidopsis genes. The latter is supported by a targeted comparative analysis between the expression profile and the chromatin folding of few marker genes (Supplemental Figure 8).

To further reveal the anchorage existing between sc/sNucRNA-seq and sNucATAC-seq experiments, we performed a correlation analysis between the expression of Arabidopsis marker genes and the folding of the chromatin at the location of their TSSs. Upon mining the scRNA-seq and sNucRNA-seq datasets, we selected the top 20 marker genes from each cluster based on their fold-change of expression compared to other clusters and p-values. Due to some redundancy between clusters, we identified a total of 378 unique marker genes. Among them, 337 are characterized by at least one sNucATAC-seq peak near their TSSs (Supplemental Table 3). As a comparison, we also performed a similar analysis on 628 Arabidopsis root housekeeping (HK) genes (i.e., genes expressed in at least in 50% of the cells and nuclei composing each cluster, across all the 20 and 21 scRNA-seq and sNucRNA-seq clusters, respectively, and with identified TSS-associated sNucATAC-seq peaks; Supplemental Table 4).

Upon normalization of their expression patterns and peak distribution across the different clusters, the marker and HK genes were distributed in 20 different bins based on their transcriptional activity and chromatin remodeling profiles. Applying the Kendall’s Tau-b rank correlation test for non-normally distributed data on the marker genes, we observed significant positive correlations between almost all the co-annotated sc/sNucRNA-seq and sNucATAC-seq datasets (Figure 4, see blue squares in the dark blue box; p-value <0.01). This result supports that the differential accessibility of the chromatin fiber correlates with the expression patterns of cluster marker genes. The similarity between the Kendall tau-b correlation maps when using the scRNA-seq and sNucRNA-seq datasets also supports the biological relevance of a single nucleus transcriptome compared to the single cell transcriptome. Based on these results, we assume that, similarly to their transcriptional activity, the remodeling of the chromatin at the location of the TSS of selected genes can be used as a molecular marker of cell type identity. As a comparison, the sc/sNucRNA-seq and sNucATAC-seq correlation analysis of the 628 HK genes only revealed few significant and more moderate correlations (Supplemental Figure 9); likely a reflection of the ubiquitous expression of these genes and the similar accessibility of the chromatin fiber near their TSS and across all the Arabidopsis root clusters.

**Figure 4.**
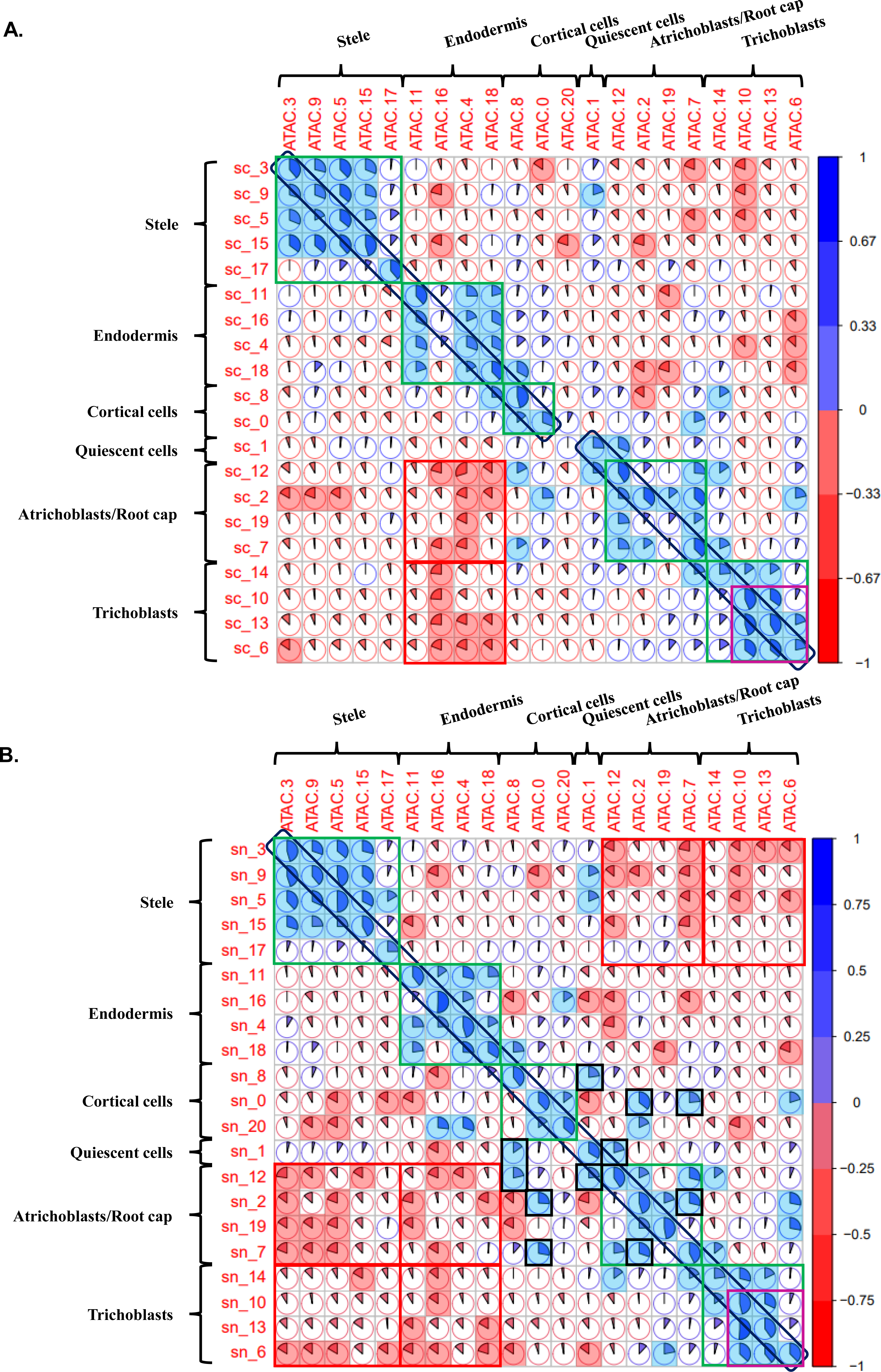
Correlation analyses between gene expression and chromatin accessibility of 337 marker genes for each of the 20 and 21 sc (**A**) and sNucRNA-seq clusters (**B**). For each correlation analysis, a Kendall tau-b correlation score was calculated based on the ranking of the cluster according to the expression level of the gene and the level of accessibility of the chromatin fiber (see pies). When significant (p-value < 0.01), positive and negative correlations are highlighted in blue or red, respectively. Co-annotated sc/sNucRNa-seq and sNucATAC-seq are highlighted in a dark purple box located in the diagonal of each figure. Positive and negative correlation between clusters that belong to super-clusters (i.e., stele, endodermis, cortical cells, quiescent cells, atrichoblasts and trichoblasts) are highlighted in green and red boxes, respectively. Other noticeable significant correlations are highlighted in black boxes.

When considering the marker genes, positive and significant correlations between sc/sNucRNA-seq and sNucATAC-seq datasets were also repetitively observed when considering the clusters that belong to the same super clusters (i.e., the “endodermis”, “stele”, “atrichoblast” and “trichoblast” superclusters; Figure 4, see blue squares in the green boxes). This result suggests that the cells composing the same tissue share, to some extent, similar transcriptomic and epigenomic signature profiles that are likely required to fulfill their tissue-specific biological functions. One exception is the cells composing the xylem (i.e., cluster #17) that seem to have unique transcriptomic and epigenomic profiles. The significant correlation existing between gene expression and chromatin remodeling suggest that the position of the nucleosomes on the double strand of the gDNA, nearby the TSS of the genes, plays a critical role in controlling the activity of the marker genes.

We also noticed positive correlations between clusters that do not belong to the same cell-types but that share similar developmental stages. For instance, the cells composting the clusters #1, 8 and 12, cells that are likely going through a differentiation process, and the cells composting the cortical and atrichoblast clusters #0, 2 and 7 share similar epigenomic and transcriptomic features (Figure 4B, see blue squares in the black boxes). These results are supported by the relative proximity of these clusters on the UMAP RNA- and ATAC-seq maps (Figure 2A and 3B). These data suggest that these cell types share similar transcriptomic programs despite their distinct ontology. In the case of the clusters#1, 8 and 12, we hypothesize that these shared programs are important in controlling plant cell differentiation and elongation processes.

### The remodeling of the chromatin at the single-cell resolution ATAC-seq can be used as a molecular marker of the root hair cell-type

Plant single cell types are annotated based on the expression profile of maker genes [7–11]. Correlation analyses between gene expression and chromatin remodeling (Figure 4) suggest that the latter could also be used as another molecular marker of the cell-types composing the Arabidopsis root. This conclusion is supported by similar analyses conducted on various animal organs and on selected plant cells [46, 52, 53]. To further test this hypothesis, we took advantage of our single cell resolution ATAC-seq datasets and focused our analysis on the three clusters representative of the mature Arabidopsis root hair cells (i.e., clusters #6, 10 and 13; Figure 2A and 5B) that share significant correlations between their transcriptome and epigenome (Figure 4, purple boxes). To validate the use of the differential chromatin accessibility as a molecular marker of plant cell identity, we first looked for the sNucATAC-seq peaks that are preferentially identified at least in two out of these three root hair clusters (fold-change of the average value across the root hair clusters versus the next highest non-root hair cluster ≥ 2). Among the 20,803 sNuc-ATAC-seq peaks, 134 were preferentially identified in the root hair mature clusters. Among them, 63 are clearly located in the TSS of annotated Arabidopsis genes. Mining the sc and sNucRNA-seq datasets, we identified 55 out of the 63 genes (87.3%) specifically expressed in Arabidopsis root hair cells independently of the transcriptomic dataset used (Figure 5; Supplemental Table 5). Among them, three genes (i.e., AT1G49100, AT1G26250, AT4G22650) were found expressed at such low level that their transcripts were detected by only sc or sNucRNA-seq technology. The remaining eight genes (i.e., AT1G07450, AT1G12050, AT2G21540, AT3G29810, AT3G53820, AT3G12530, AT5G39100, and AT5G41290) were not found as preferentially expressed in root hair cells. Initiating our analysis by identifying the peaks of accessible chromatin specific to the clusters #6, 10, and 13 led to the identification of associated root hair maker genes. In addition to support the role of chromatin remodeling in controlling gene expression in plant cells, this work highlights that the remodeling of the chromatin can be used as a molecular marker to annotate specific cell types.

**Figure 5.**
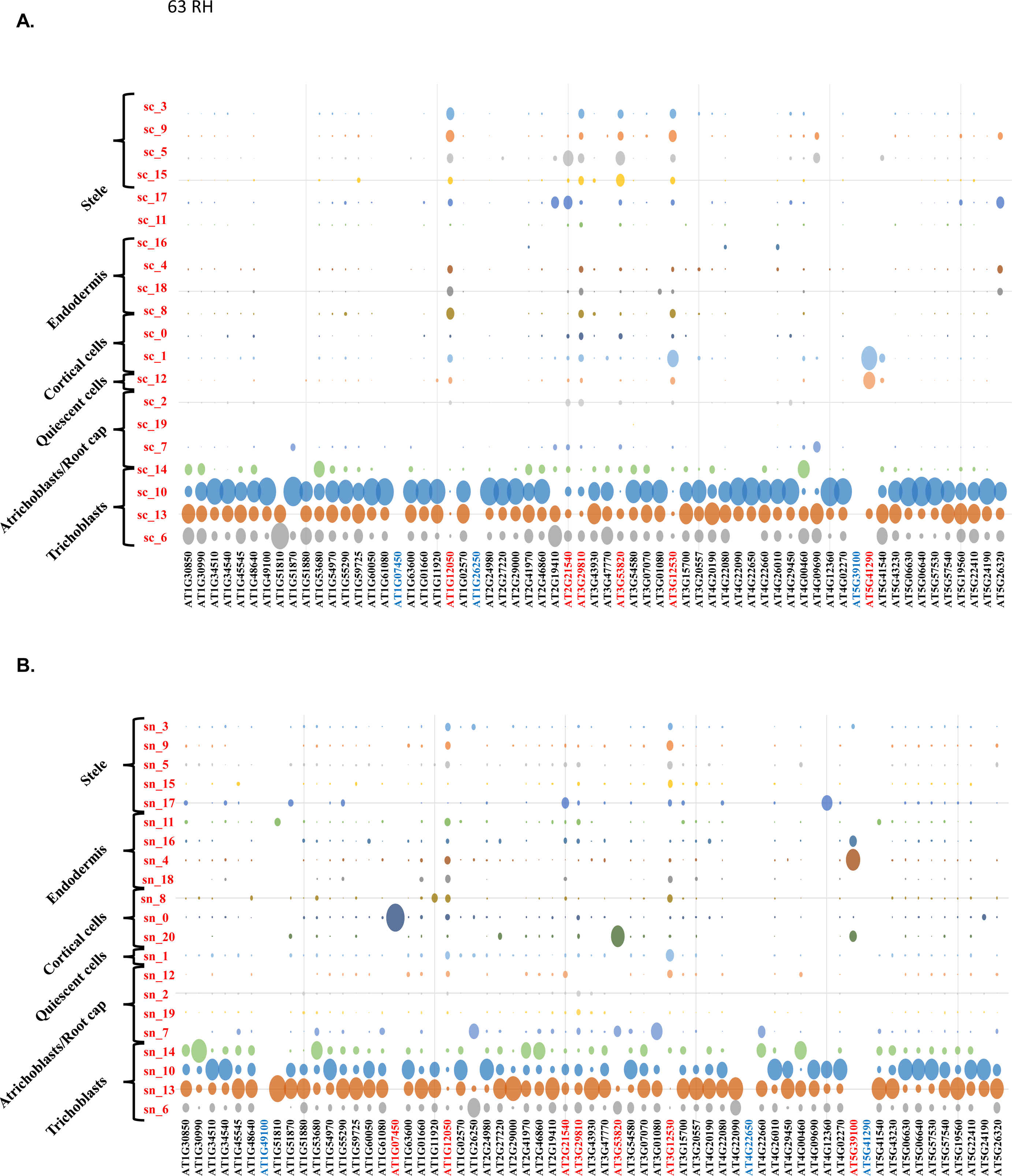
Normalized expression level of 63 Arabidopsis genes (x-axis) characterized by their higher accessibility of their TSS in the root hair clusters #6, 10, and 13 (Figure 4B). Gene expression levels were quantified by using scRNA-seq (A) and sNucRNA-seq (B). The diameter of each circle reflects the relative expression of the genes in each cell cluster (y-axis). Genes highlighted in red are not considered as root-hair specific based on their transcriptional patterns. Those highlighted in blue are not found expressed according to the sc (A) and sNucRNA-seq datasets (B).

## Conclusions

In this manuscript, we describe the use of isolated nuclei as an alternative approach to the use of plant protoplasts to obtain biologically informative transcriptomic information at the single cell level. Starting with a limited pool of poly-adenylated transcripts, the number of transcripts sequenced and, consequently, the number of expressed genes per nucleus are lower compared to the protoplast-based approach (Supplemental Table 1). However, we found that the nuclear transcriptome strongly correlates with the cellular transcriptome (Figure 2). Unexpectedly, the sNucRNA-seq technology complements the published protoplast-based Arabidopsis root transcriptome by defining one new cell clusters (i.e., cluster #20), and by substantially enriching for two additional clusters (i.e., clusters #16 and 19).

We also describe a global map of chromatin accessibility of the Arabidopsis root at a single cell-resolution. This level of resolution revealed the unique and conserved usage of the genomic information between the cells. Assuming that cluster-specific remodeling of the chromatin could be associated with a cluster-specific transcriptome, and, therefore, with a specific Arabidopsis root cell type, we performed a multimodal integration between chromatin accessibility and transcriptomic profiles using a set of cell-type marker genes. Independently of the use of scRNA-seq or sNucRNA-seq, we were able to highlight significant correlations between the two types of datasets for co-annotated clusters. Interestingly, a significant correlation exists between clusters that are not co-annotated but that belong to the same super-clusters (Figure 4). Together, our data support a role for chromatin accessibility in controlling gene expression. They also demonstrate that many marker genes exhibit cell-type specific remodeling of the chromatin. To further explore this, we performed a reverse analysis by first identifying accessible chromatin near TSSs and then mining our sc/sNucRNA-seq dataset. In this way, we were able to confirm that open chromatin near the TSS correlates well with transcriptionally active genes in specific cell clusters. Thus, our work reveals that the chromatin accessibility could be used as another molecular marker of plant cell types.

The use of nuclei to access both transcriptomic and epigenomic datasets can be further explored by analyzing the role of epigenomic marks in gene expression. Expanding the portfolio of single nuclei –omics technologies would help understand the influence of epigenome and epigenomic marks on gene expression. In addition, multiomic approaches at the level of single nuclei could enhance comparative analysis between plant species. The use of isolated nuclei avoids many of the technical and biological difficulties associated with protoplasts and so will likely enable the broad analysis of gene expression across different species and tissues.

## Methods

### Plant growth

Arabidopsis (*Arabidopsis thaliana,* Col-0 ecotype) seeds were surface sterilized using ethanol 70% then a 30% (v/v) bleach, 0.1% (v/v) Triton X-100 solution for 5 min each. The seeds were then placed on Murashige and Skoog (MS) growth media agar plate under 16 light and 8 dark conditions. Seven days after germination, the primary roots of around 150 seedlings were isolated from the seedlings.

### Root Nuclei Isolation and single nuclei RNA-seq and ATAC-seq library constructions

To maximize the relevant of the transcriptomic information collected from plant isolated nuclei, we developed a methodology to isolate nuclei and process them for reverse transcription. The release of the plan nuclei is accomplished in no more than 5 minutes and exclusively occurs at 4°C to minimize RNA degradation, prevent nuclear membranes, and prevent transcriptional changes associated with the isolation process. As a comparison, 60 to 120 minutes incubation at room temperature to 25°C are required to digest the cell wall to release Arabidopsis root protoplasts [7–11]. Upon centrifugation, FANS, and estimation of the nucleate density, the nuclei are processed into the 10X Genomic Chromium System for the construction of the sNUC-RNAseq libraries in less than 90 minutes following the isolation of the plants from the Petri dishes.

To generate sNucRNA-seq libraries, Arabidopsis root nuclei were isolated by chopping the roots in 1 ml of Nuclear Isolation Buffer (CELLYTPN1-1KT, Sigma-Aldrich) and incubated for 15 minutes on a rocking shaker at 4°C. The lysate was filtered through a 40 μm cell strainer (Corning) and stained with propidium iodide (PI; 20 ug/ml, Alpha Aesar). Nuclei were sorted using FACSAria™ II flow cytometer (BD, Flow Cytometry Service Center), pelleted (1,000g for 10 minutes at 4°C), and resuspended in PBS buffer without calcium and magnesium and supplemented with Protector RNase inhibitor (0.2U/ul, Roche) and ultrapure BSA (0.1%, Invitrogen). Nuclei concentration was estimated using Countess II FL Automated Cell Counter (ThermoFisher) upon staining with trypan blue. The nuclei were used as an input to generate RNA-seq libraries following the 10X Genomics Chromium Single Cell 3□ v3 protocol without modification.

To create single nuclei ATAC-seq libraries, the nuclei were isolated by chopping and filtration as described above. Triton X-100 (v/v) 0.5% final was added to the nuclei suspension to permeabilize the nuclei. The nuclei were then centrifuged 500g for 10 minutes, washed with PBS supplemented with BSA 0.1%, then pelleted again. Oppositely, to the population of nuclei used to establish sNucRNA-seq libraries, we did not stain with propidium iodide and sorted the nuclei dedicated to sNucATAC-seq technology. Preliminary work conducted on Arabidopsis nuclei clearly highlighted that both staining and sorting decrease the number of unique fragments per barcode, affecting the resolution of the clusters of cells (data not shown). Upon isolation and filtration, the nuclei pellet was resuspended in the ATAC-seq nuclei buffer (10x Genomics #2000153). Nuclei concentration was estimated using the Countess II FL Automated Cell Counter (ThermoFisher). 6768 nuclei isolated from two independent replicates were used to generate sNucATAC-seq libraries following 10X Genomics Chromium Single Cell ATAC protocol without modification.

The sNucRNA-seq and sNucATAC-seq libraries were sequenced on an Illumina Hiseq platform.

### Bioinformatics analyses

*Sc/sNucRNA-seq:* ScRNA-seq data for the three wild-type replicates of previously published [9] data (SRA experiments SRX5074330-SRX5074332) and five sNucRNA-seq replicates from the current study (Supplemental Table 1) were processed individually using the 10x Genomics cellranger (v3.1.0) count pipeline, against a reference constructed from the TAIR10 genome and Araport11 annotations. The recommended protocol for reference construction in support of pre-mRNA capture for data derived from nuclear preparations was applied.

Following generation of the UMI count matrices, doublet filtering was performed on each individual dataset using 50 iterations of the BoostClassifier prediction method of DoubletDetection (p_thresh=1e-7, voter_thresh=0.8) [54]. Additionally, the published scRNA-seq data was filtered to include only those cells passing QC as performed in [9] by filtering on cell barcodes present for the WT replicates in the GEO dataset GSE123013. All datasets were filtered to include only called cells with more than 200 genes represented by at least one UMI. Normalization of individual datasets and their integration was performed using Seurat v3.1.5, with genes required to be present in a minimum of 5 cells, and the top 2000 variable genes used for feature selection. Integration anchors were chosen for the combined set of 3 scRNA-seq and 5 sNucRNA-seq replicates using the first 20 dimensions of the canonical correlation analysis method. Following integration, UMAP dimensionality reduction was performed on the first 20 principal components, and clustering was performed using Seurat’s FindClusters method with a resolution of 0.5. Expression values were obtained separately for the subsets of cells and nuclei belonging to each cluster using Seurat’s AverageExpression method.

#### SNucATAC-seq

SNucATAC-seq data from 2 replicates were processed using the cellranger-atac (v1.2) count pipeline, with a reference constructed using TAIR10/Araport11 and JASPAR2018_CORE_plants_non-redundant_pfms. Counts against peaks called by cellranger-atac were read into Seurat objects and subset to include only cells with between 1k-20k fragment counts in peak regions, and > %15 of the cell’s fragments in peak regions and a calculated nucleosome_signal of > 10. Integration of the two replicates was performed using the Harmony integration strategy [55] on the latent semantic indexing reduction performed by Signac (v0.2.5) [51], with UMAP dimensionality reduction performed on the resulting integrated dataset. Transcription start site (TSS) enrichment was analyzed per chromosome using the TSSEnrichment/TSSPlot functions of Signac.

### UMAP visualization

In order to relate the structure of the UMAP plot for the combined sc/sNucRNA-seq datasets to that of the sNucATAC-seq data, a putative gene activity matrix was constructed from the aggregated fragments of the two replicates, using gene body coordinates extended 2kb upstream of the transcription start site. Log normalization was performed on the activity matrix, then the Seurat FindTransferAnchors method was used with the sNucATAC-seq data as query against the combined RNA-seq data as reference. Finally, cluster assignments were made on the sNucATAC-seq using the TransferData method and the integration anchors determined earlier. These cluster assignments were used for subsequent extraction of averaged fragment counts across all cells assigned to the cluster, using Seurat AverageExpression on the peaks.

For visualization purposes, data from all RNA-seq data were combined using cellranger aggregate method to produce a combined cloupe file, with UMAP coordinates and cluster assignments imported from the Seurat analysis. Similarly, the two sNucATAC-seq replicates were combined using cellranger-atac aggregated, and UMAP coordinates and cluster assignments from the Harmony/Signac analysis were imported into the cloupe rendering.

#### Correlation analyses

Correlation analyses were performed by matching called peaks from the ATAC-seq data to genes by looking for any overlap of peak regions with genes using bedtools (v2.29.2) [56], then applying a custom script to choose among overlapped genes based on the maximum extension of the peak region into the 5’ upstream region. Correlation was performed between matrices constructed from normalized average expression values for genes and average fragment count values for peaks, with the corresponding rows in the two matrices representing matched gene and peak entities. The normalization procedure was done on values in each row independently, by considering each cluster in terms of its percentage representation of the expression/fragment count for the gene/peak across all clusters, in order to represent specificity independent of magnitude of expression/fragment count. These normalized values were considered as mapping onto twenty quantiles, roughly corresponding to the number of clusters in the analysis. Thus, a gene with perfect specificity would be mapped into the highest quantile for its cluster context of expression and into the lowest quantile for all other clusters, while a gene expressed approximately equally across all clusters would be mapped into the next-to-least quantile for each cluster. The R cor function with method=”kendall” (implementing the Kendall tau-b method for rank-based correlation in the presence of tied ranks) was used to produce correlation matrices, which were visualized as correlograms from the corrplot package [57].

## Data Availability

Expression data are available at the Gene Expression Omnibus (GEO number: pending).

## Supporting information

Summary of the sequencing of the different single cell and single nuclei RNA-seq libraries.

List of Arabidopsis root cell-type marker genes.

Expression level and relative accessibility of the chromatin of 337 Arabidopsis root cell-type marker genes.

Expression level and relative accessibility of the chromatin of 628 Arabidopsis root housekeeping genes.

Expression level and relative accessibility of the chromatin of 63 Arabidopsis genes associated with root hair-preferential sNucATAC-seq peaks in thei

## Acknowledgements

This work was supported by a grant to M.L. from the U.S. National Science Foundation (IOS# 1854326), by grants to J.S. from the U.S. National Science Foundation (IOS#1923589) and the Department of Energy (DE-SC0020358), by the Center for Plant Science Innovation, and by the Department of Agronomy and Horticulture at the University of Nebraska-Lincoln. The authors would like to acknowledge Dirk Anderson, manager of the Flow Cytometry Core facility, and the Nebraska Center for Biotechnology at the University of Nebraska-Lincoln, for providing support in the sorting of the isolated nuclei. Additionally, the authors want to thank Dr. Jeffrey Doyle, from Cornell University for his constructive comments and feedback on the manuscript.

## Author Contributions

S.T. and K.H.R. performed experiments. A.D.F., S.C., K.H.R., J.S., and M.L. carried out data analysis. M.L. oversaw the study. All authors contributed to the preparation of the manuscript.

## Legends

**Supplemental Figure 1.**
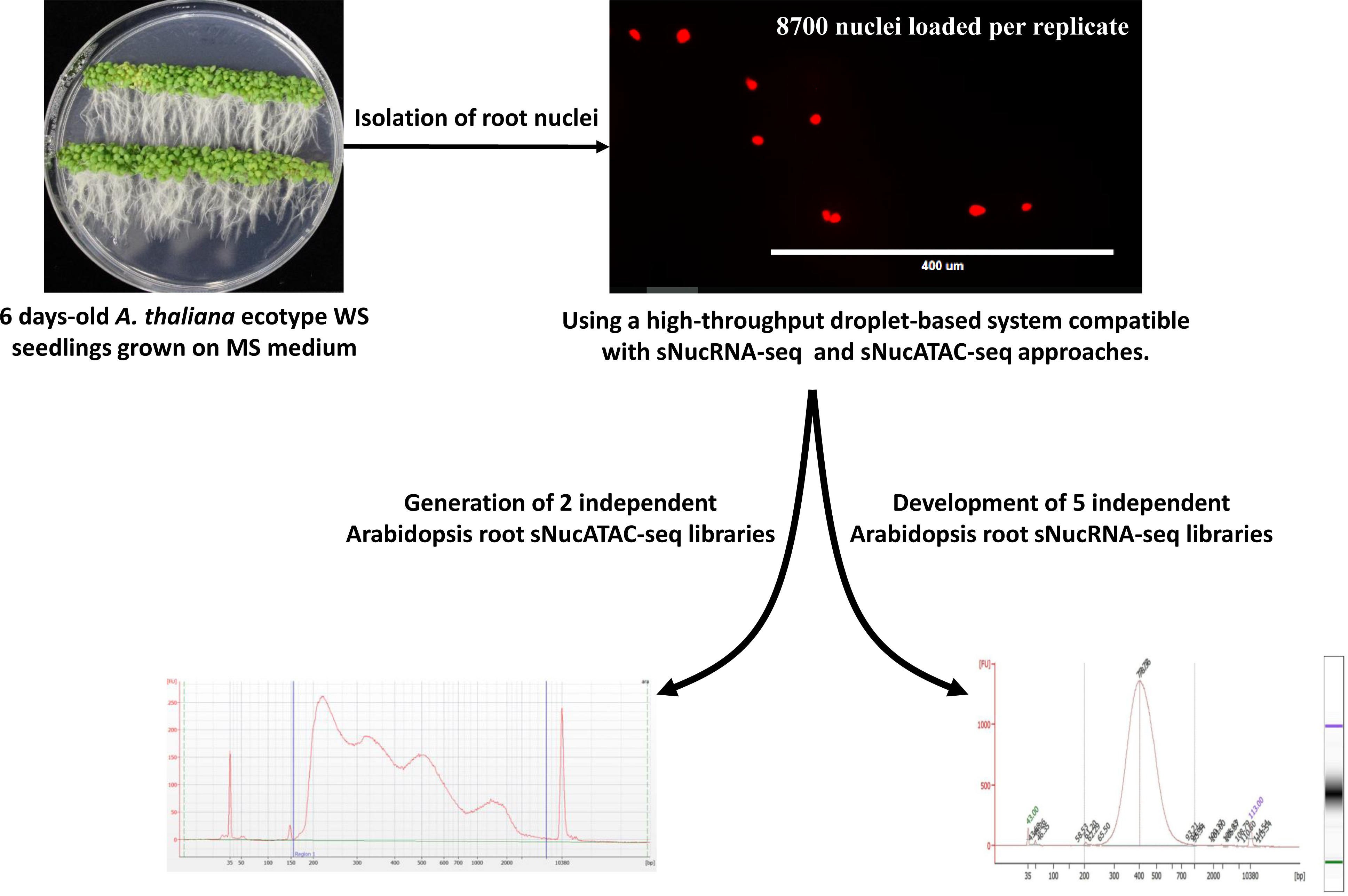
Schematic representation of the isolation of Arabidopsis root nuclei and their use to create sNucRNA-seq and sNucATAC-seq libraries.

**Supplemental Figure 2.**
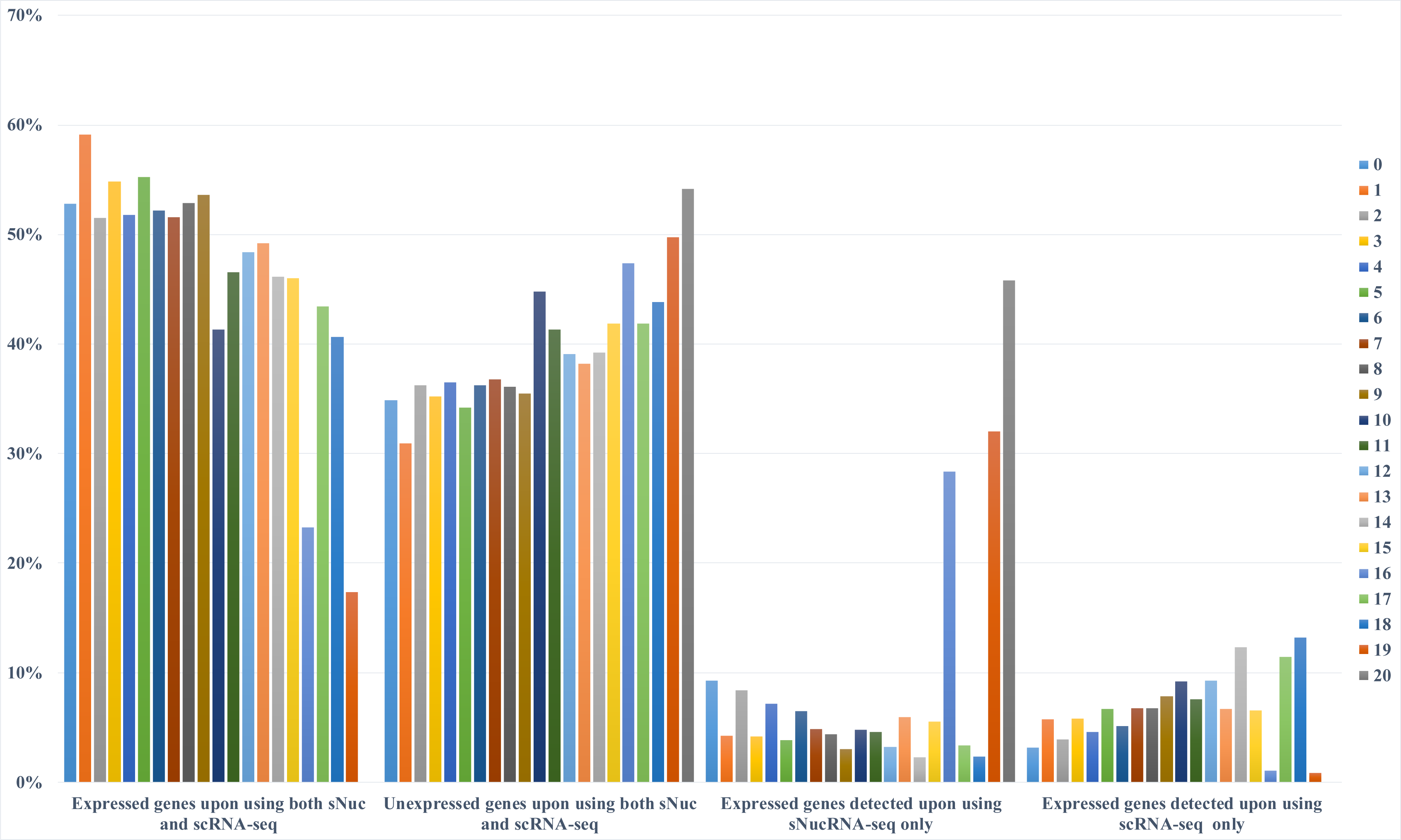
Percentage of the predicted Arabidopsis protein-coding genes found expressed or unexpressed by using sNucRNA-seq and scRNA-seq technologies. These percentages are provided for each of the 21 clusters described in Figure 1A and B. As a note, the transcriptomes of three clusters (clusters #16, 19, and 20) almost exclusively relay on the use of sNucRNA-seq technology.

**Supplemental Figure 3.**
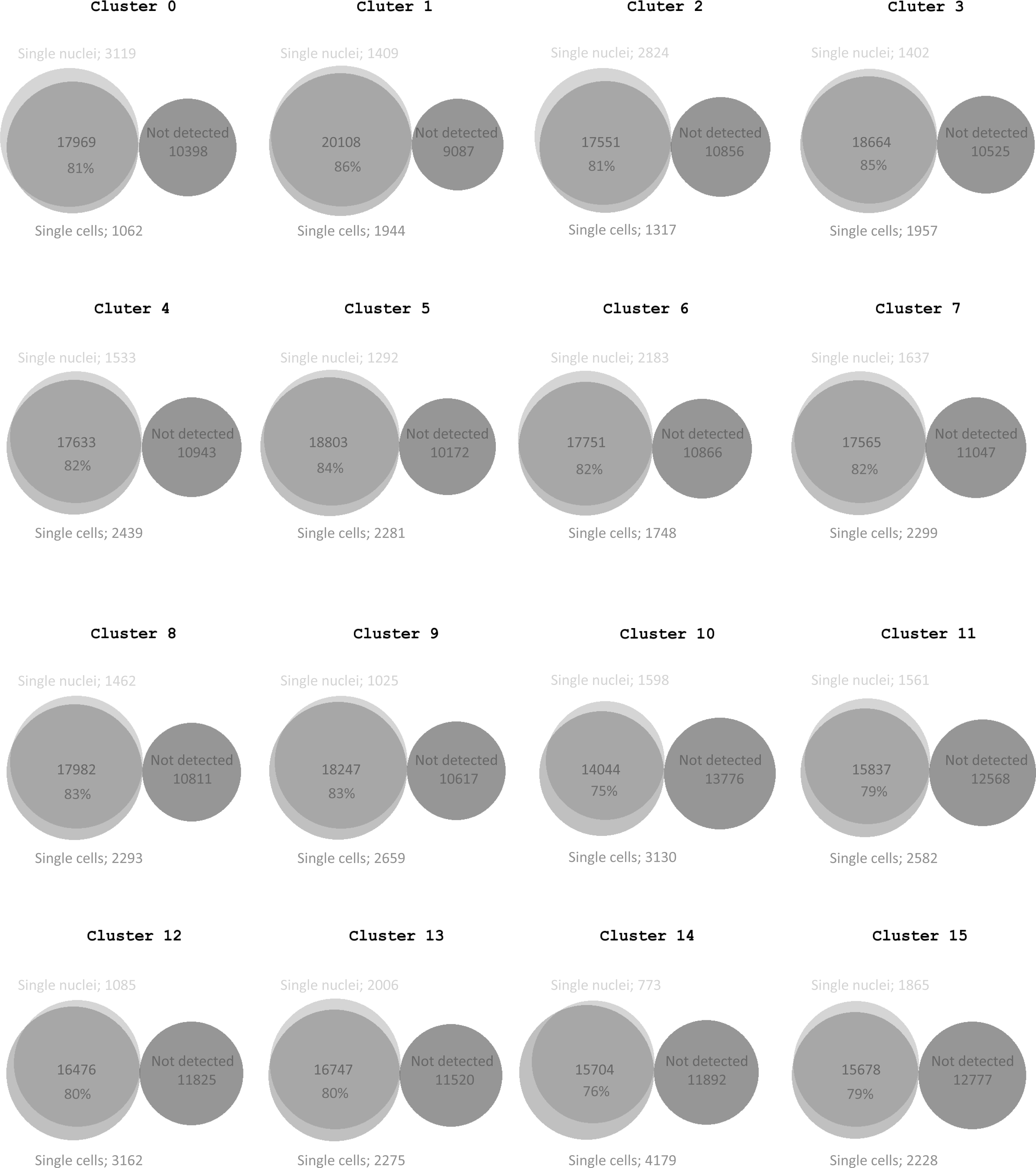
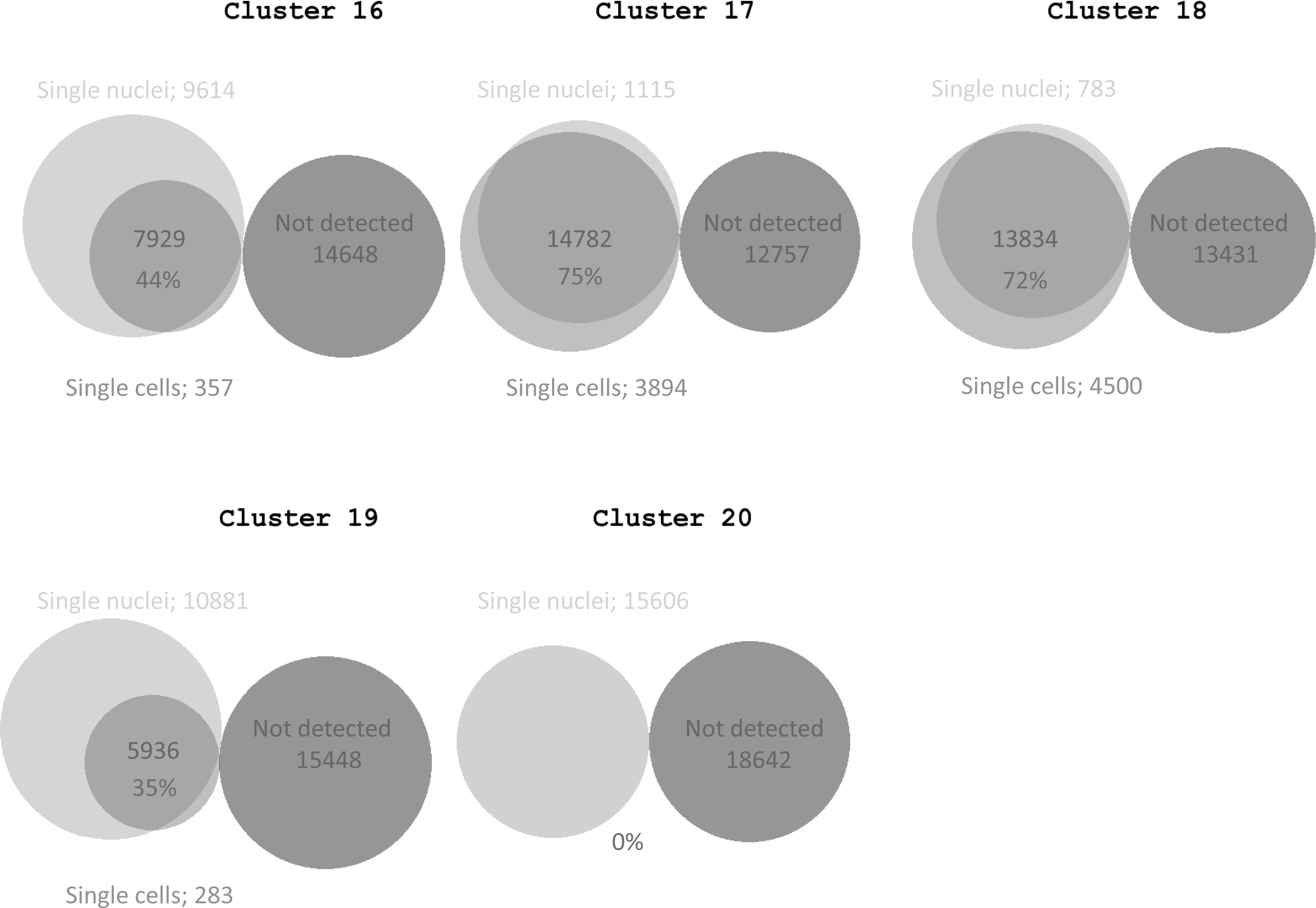
Venn diagrams showing the number of expressed and non-expressed genes revealed upon applying sNucRNA-seq and scRNA-seq technologies. Not including clusters #16, 19 and 20, 80.2% of the expressed genes were identified by both technologies in average.

**Supplemental Figure 4.**
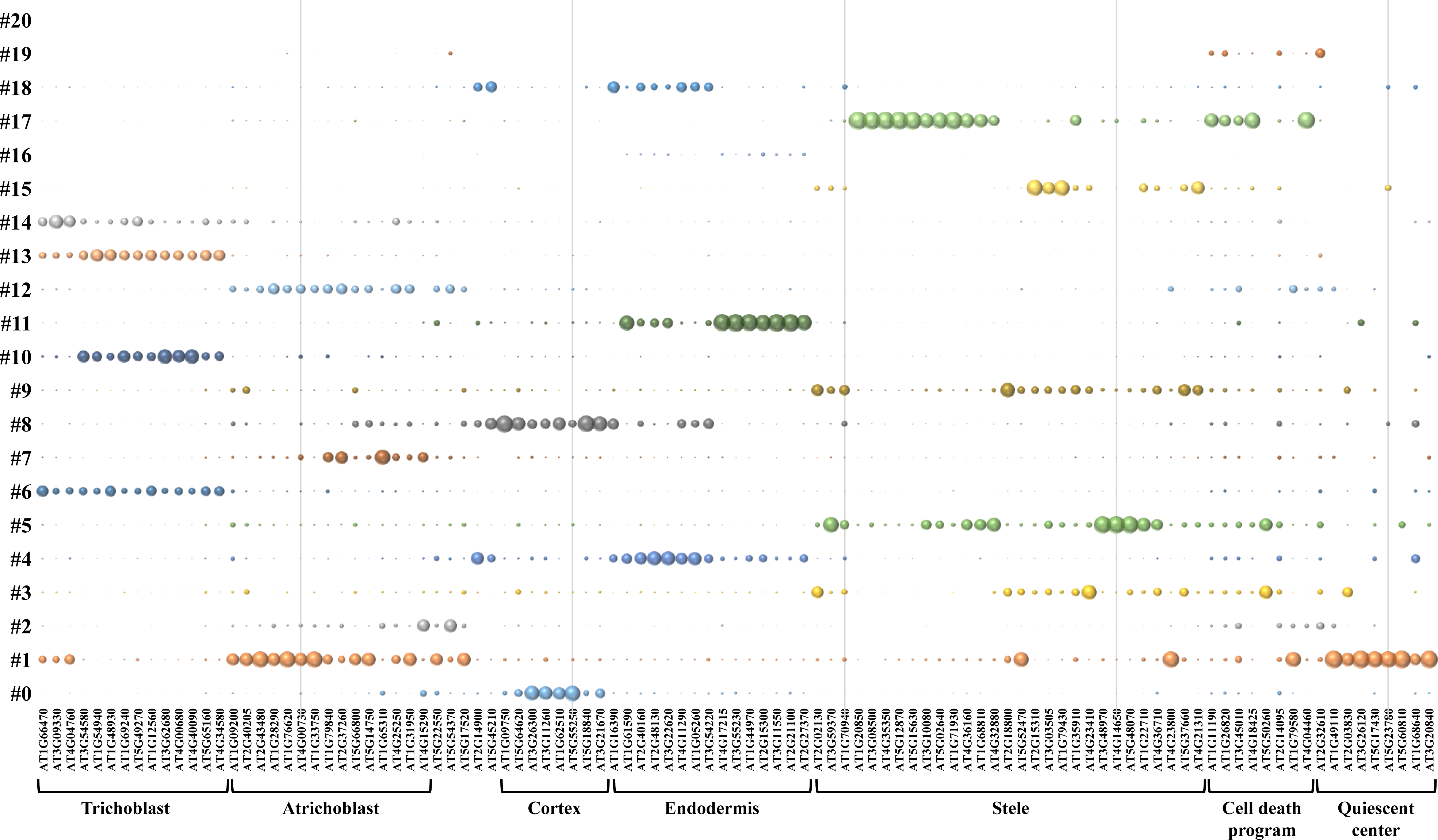
Normalized expression level of 103 cell-type marker genes (x-axis) across the 21 different clusters (y-axis). scRNA-seq dataset mined from Ryu et al., 2019 [9] were used to create this figure. The diameter of each circle reflects the relative expression of the genes in each cell cluster.

**Supplemental Figure 5.**
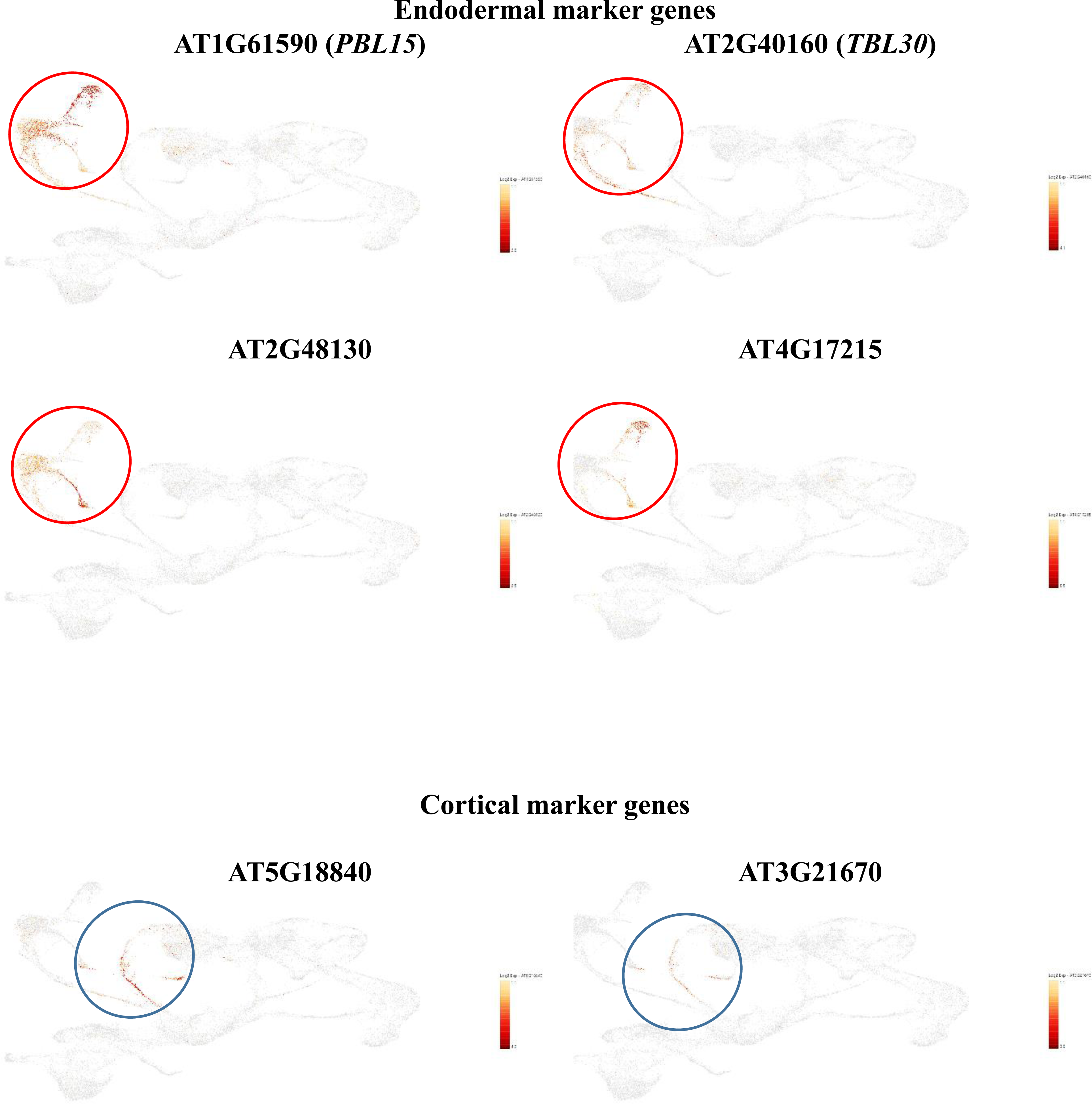
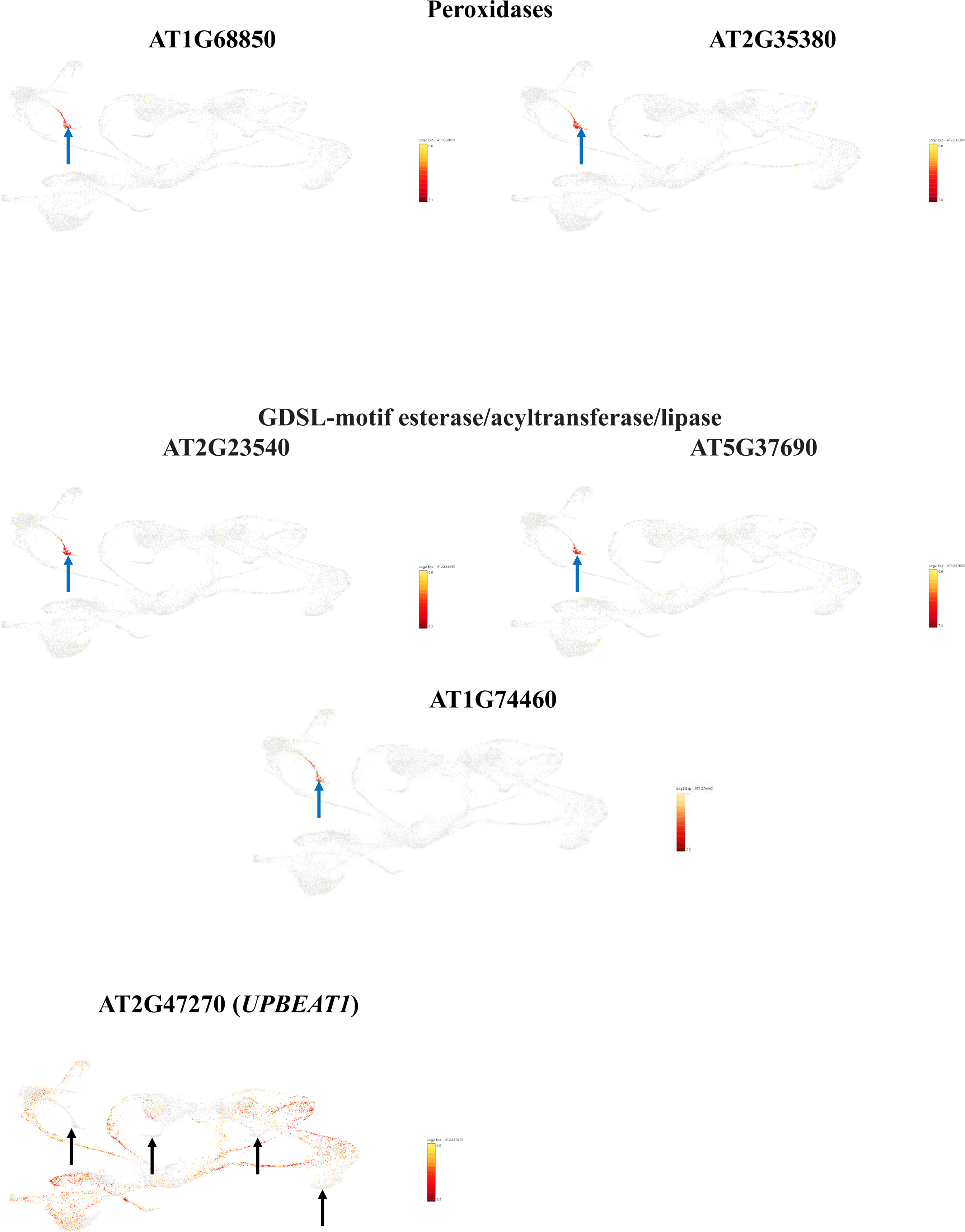
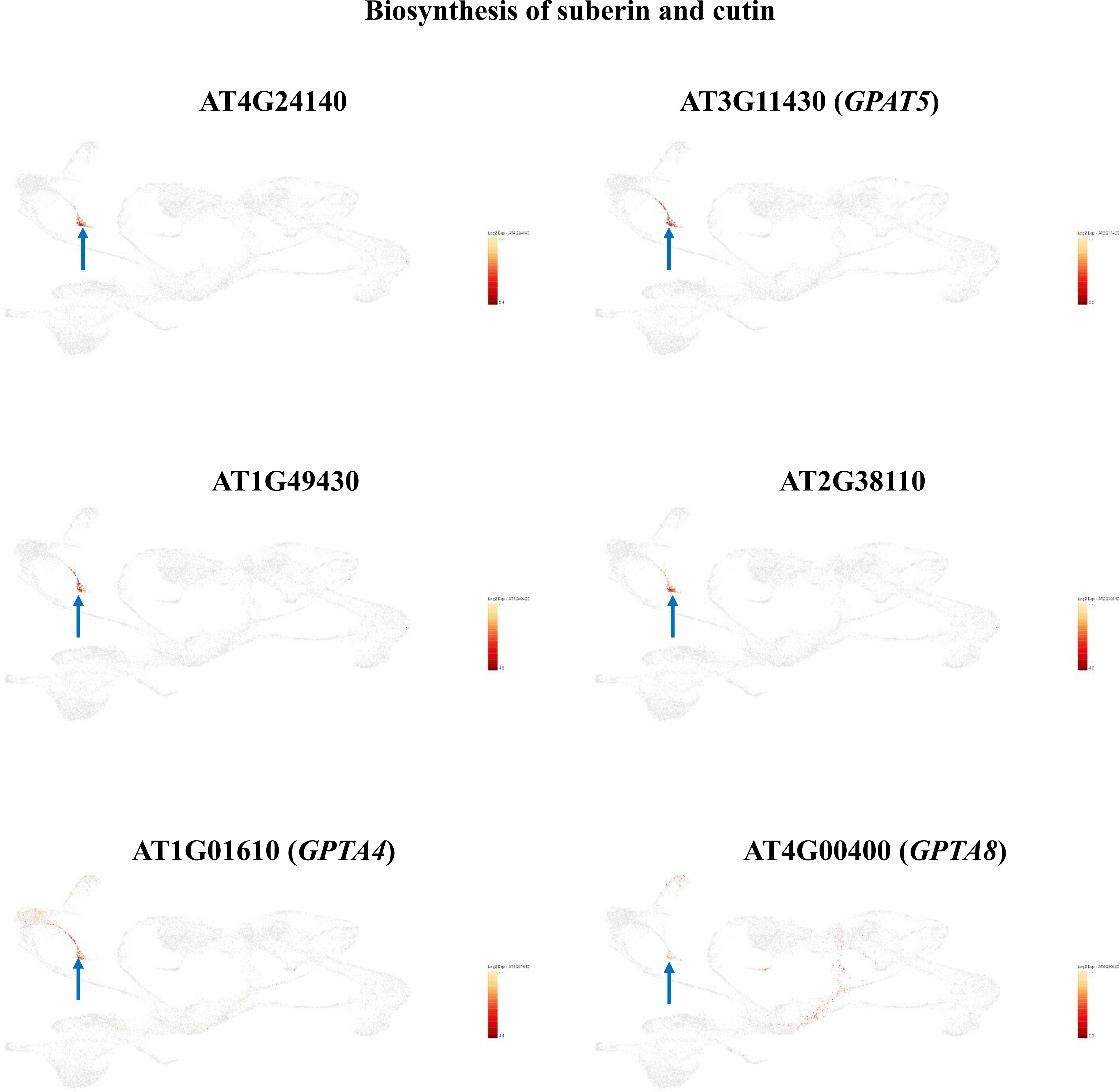
Transcriptional patterns at the single cell level of Arabidopsis genes preferentially expressed in cluster #16. The relative levels of expression of the genes are highlighted in yellow/red color. Red and blue circles highlight the expression of endodermal and cortical marker genes. Blues arrows highlight the expression of gene encoding peroxydases, GDSL-motif esterases/acyltransferases/lipases and proteins involved in the biosynthesis of suberin and cutin. Black arrows highlight the absence of transcriptional activity of the *UPBEAT1* gene in the clusters # 10, 16, 19 and 20.

**Supplemental Figure 6.**
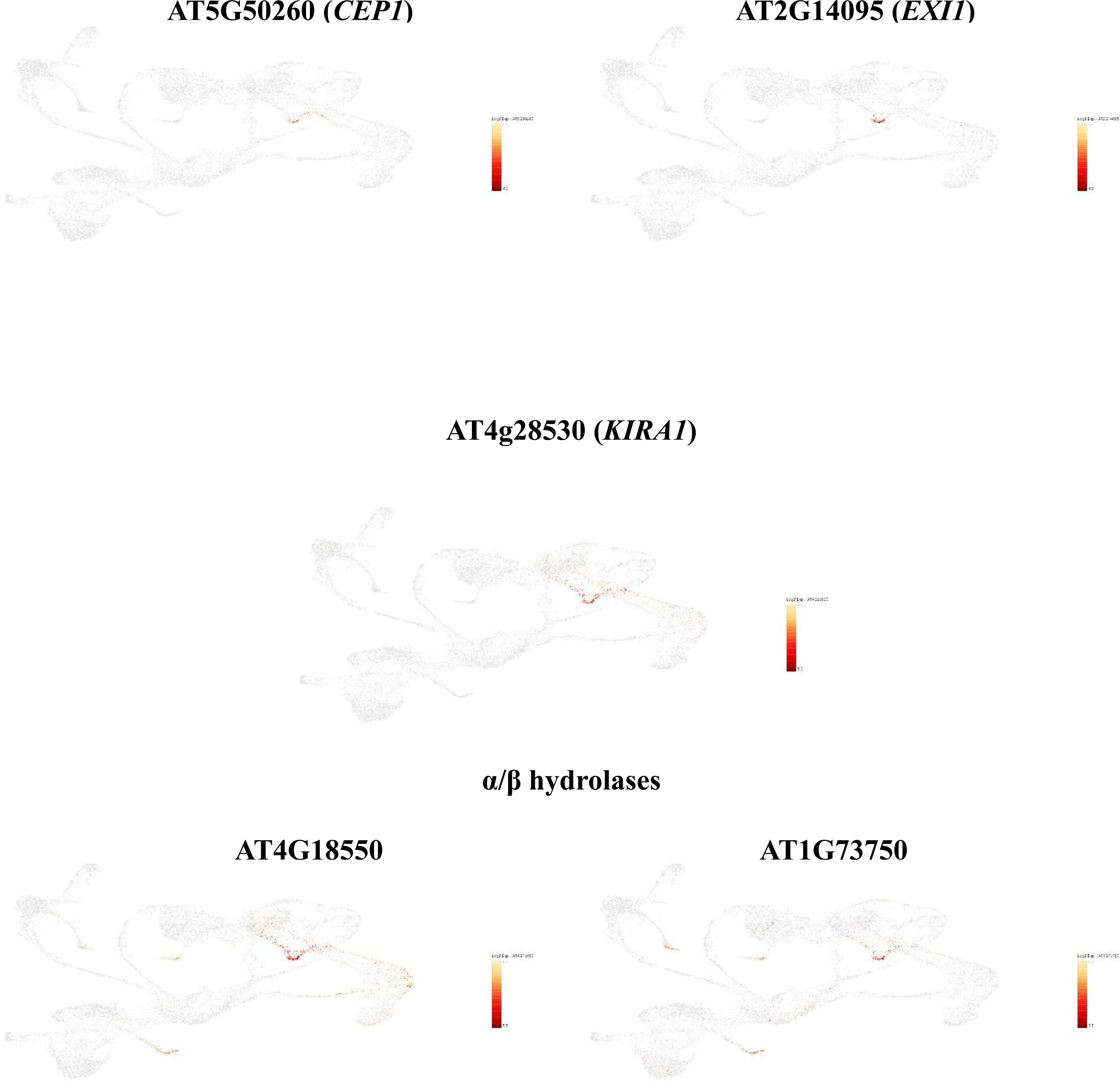
Transcriptional patterns of the Arabidopsis *CEP1* (AT5G50260), *EXI1* (AT2G14095), and *KIRA1* (AT4G28530) genes in isolated Arabidopsis nuclei (B). These genes control the cell death program notably in the root cap. The expression profile of two α/β hydrolases potentially involved in suberin/cutin biosynthesis is also highlighted. The relative levels of expression of the genes are highlighted in yellow/red color.

**Supplemental Figure 7.**
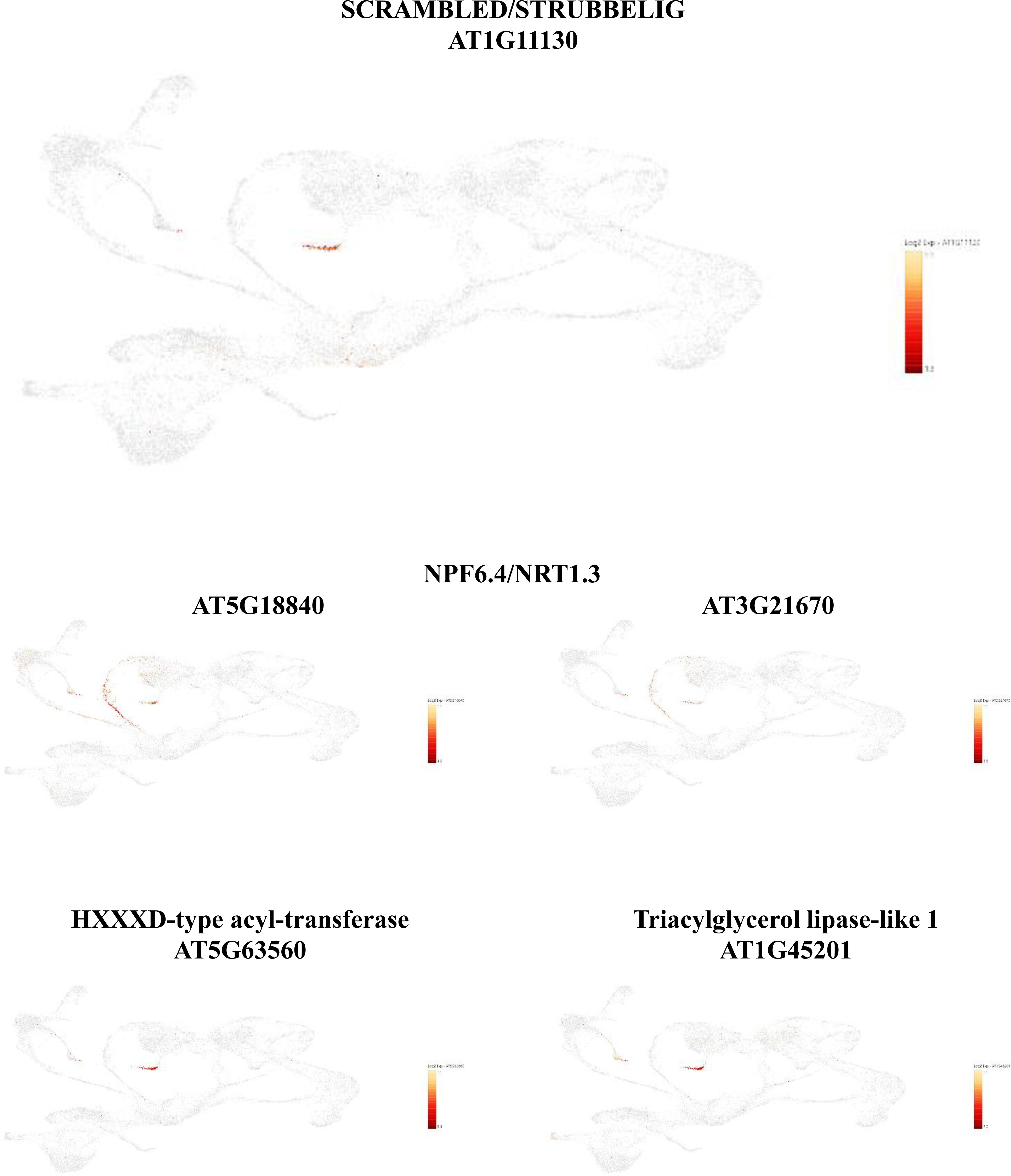
Transcriptional patterns of the Arabidopsis *SCRAMBLED/STRUBBELIG* (SCM, AT1G11130) gene, two *NPF6.4/NRT1.3* genes, and couple additional marker genes of this cluster playing a role in root hair cell differentiation. The relative levels of expression of the genes in isolated Arabidopsis nuclei are highlighted in yellow/red color. SCM was mostly expressed in cluster #20, a cell cluster exclusively identified by applying sNucRNA-seq technology. In protoplast, the expression levels of these genes are limited to few cells.

**Supplemental Figure 8.**
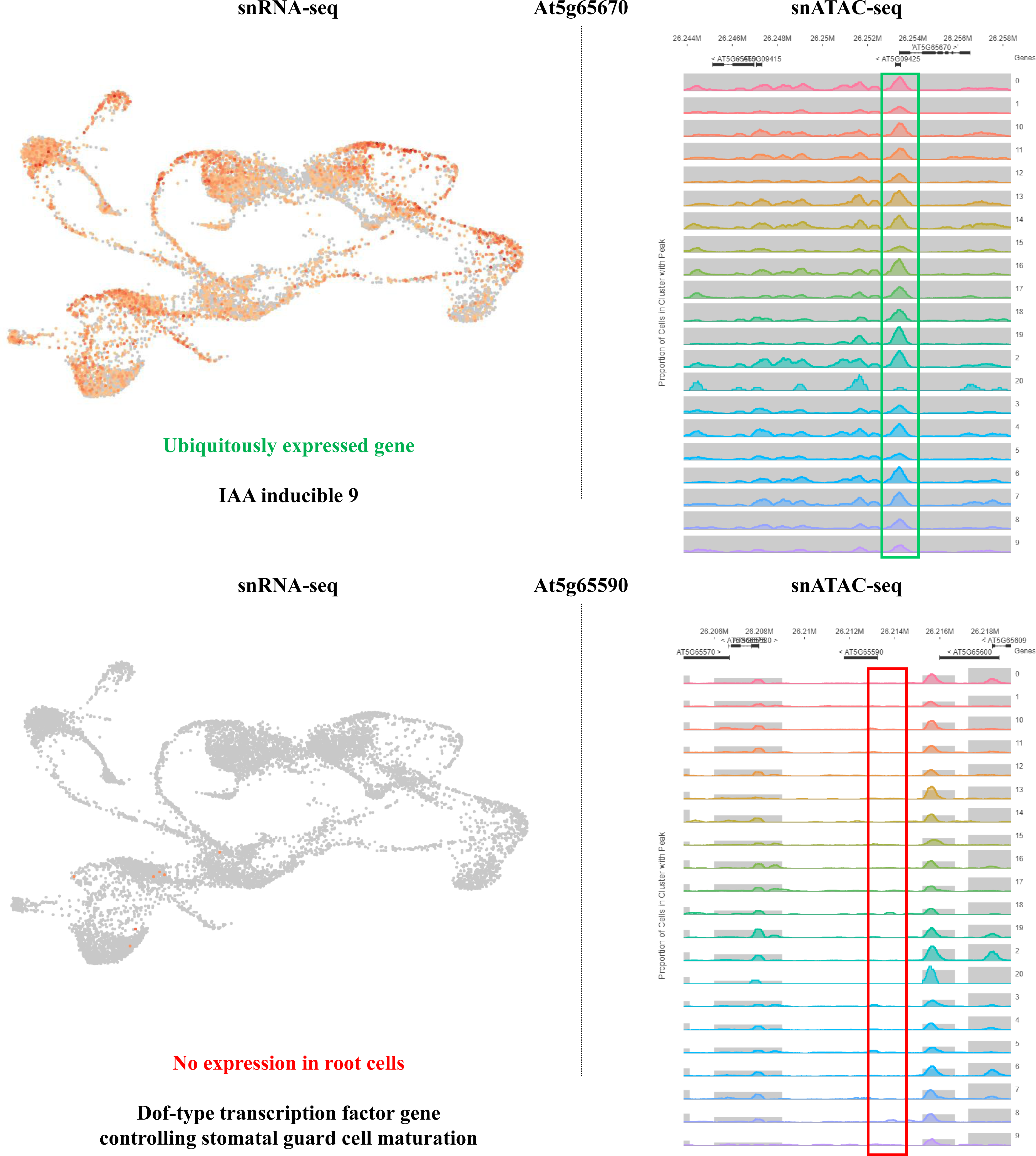
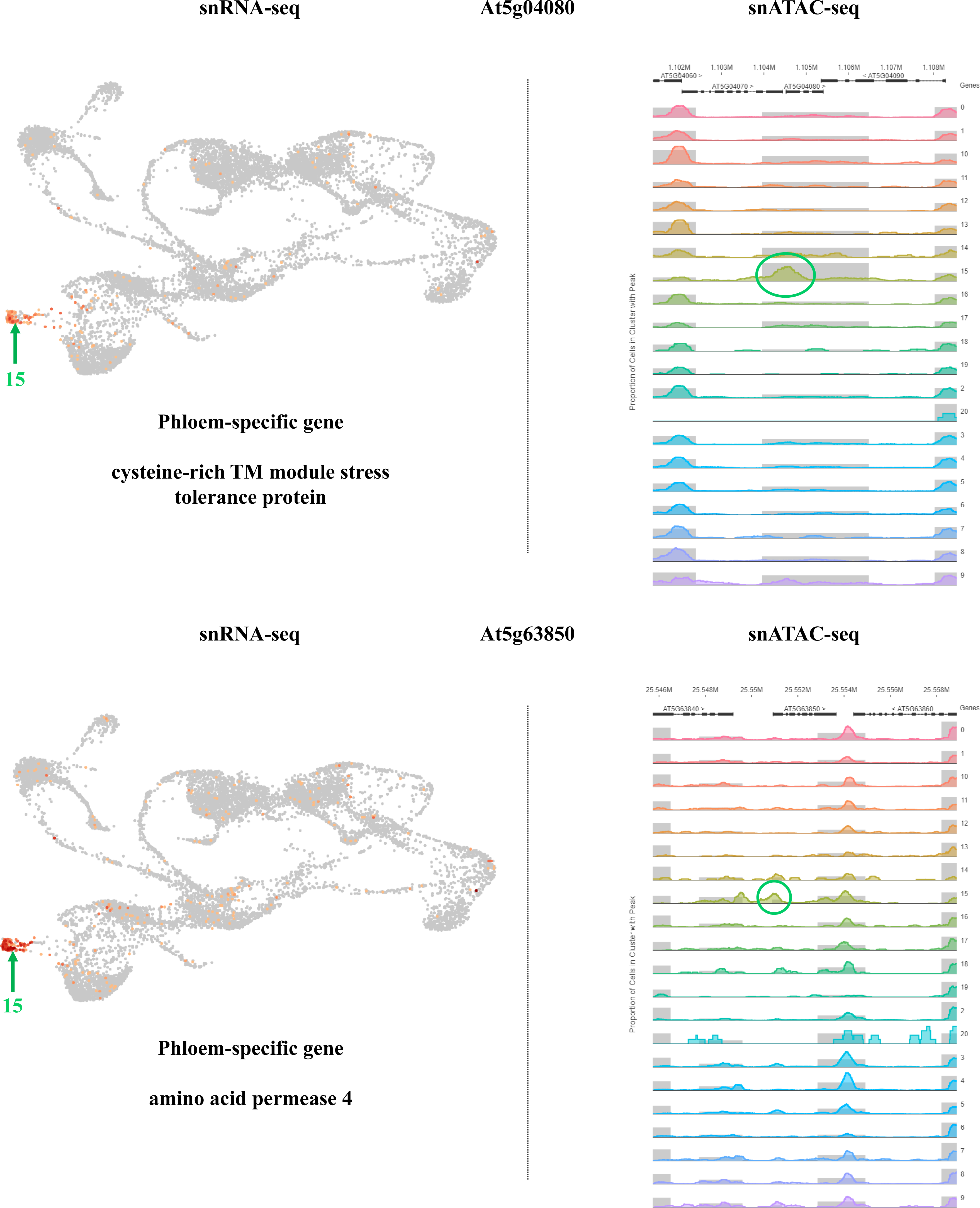
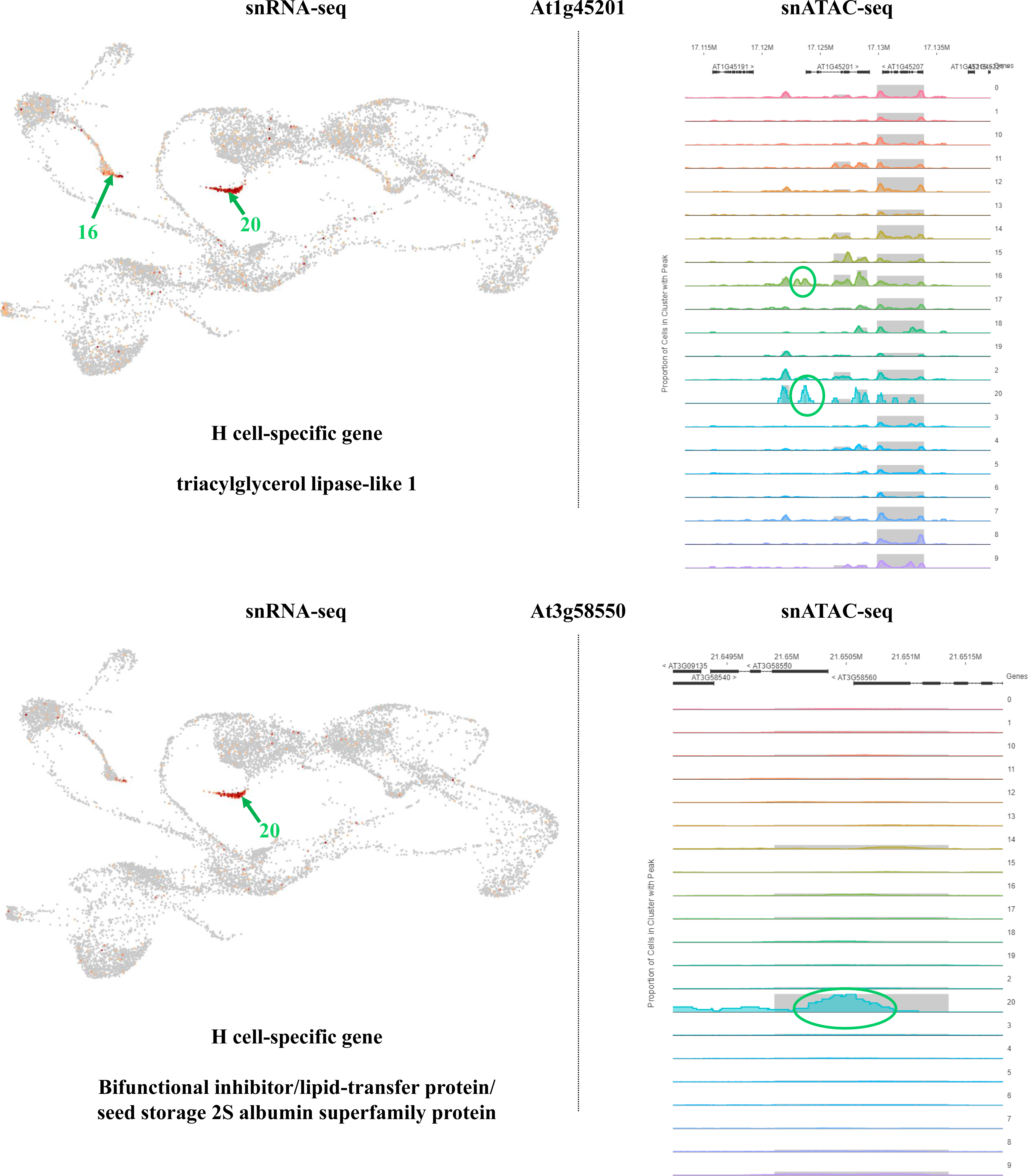
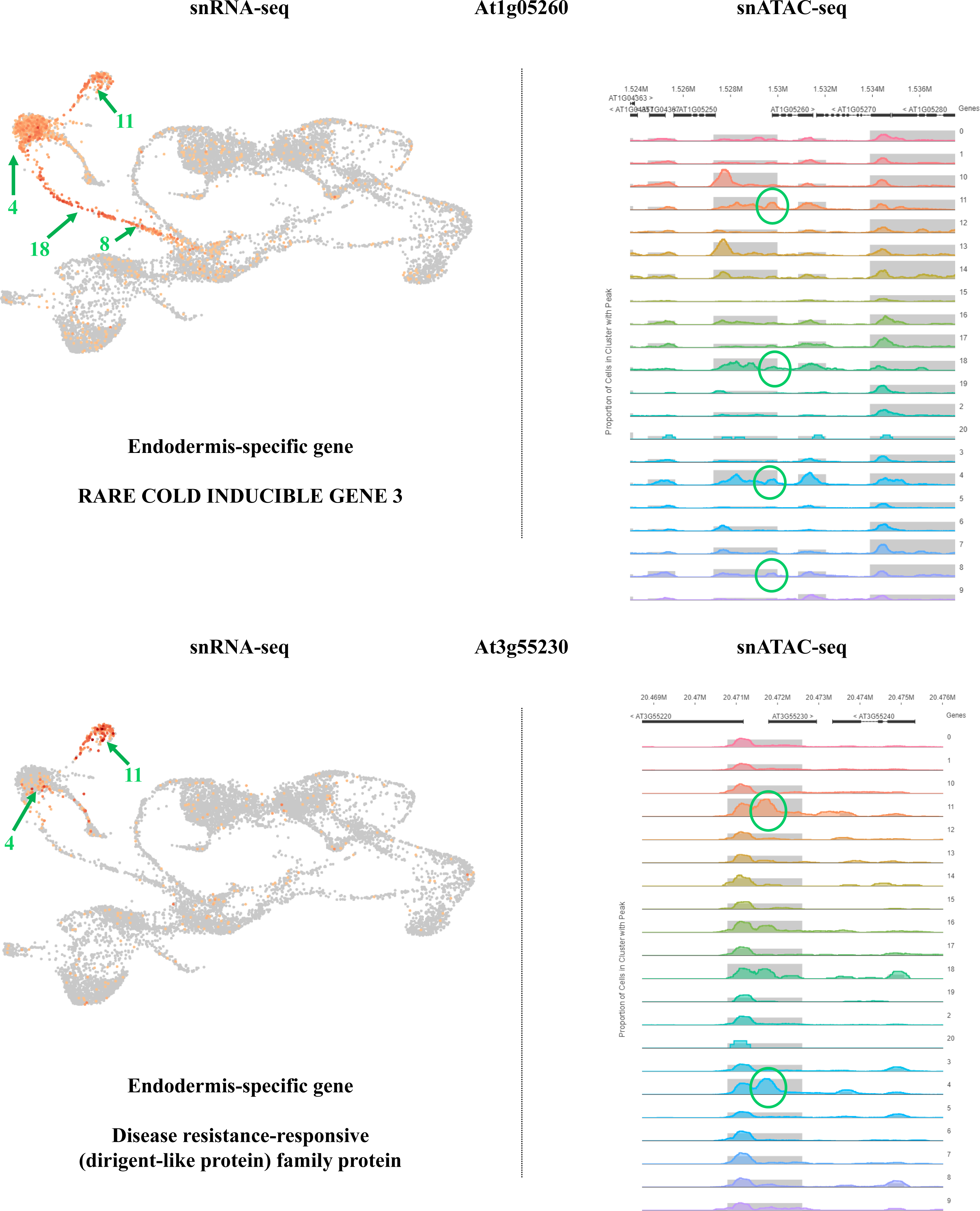
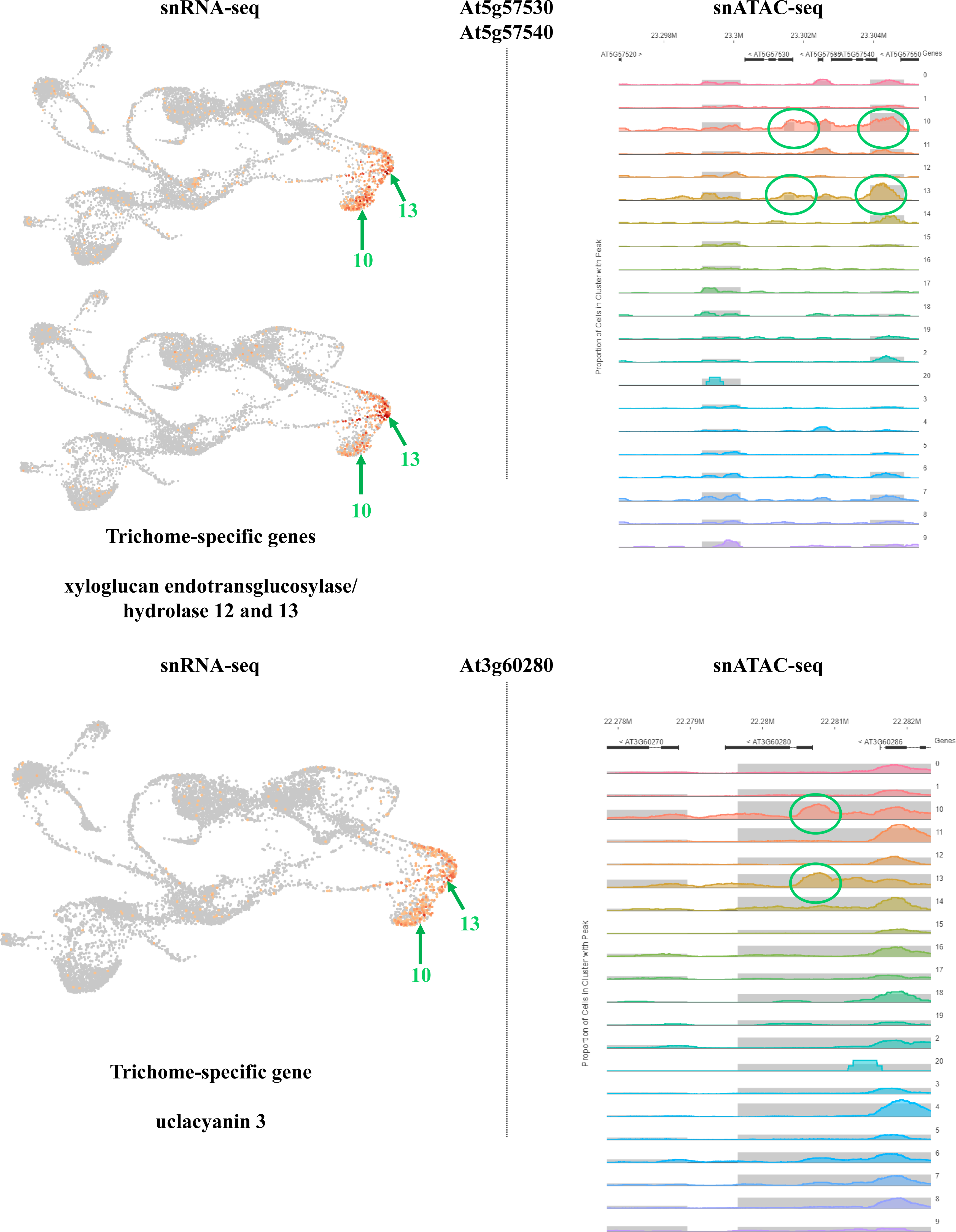
Comparative analysis of the expression profile and chromatin folding of selected Arabidopsis genes. The left panels highlight the transcriptional activity of selected genes according to sc/sNucRNA-seq datasets (the relative levels of expression of the genes are highlighted in yellow/red color). When relevant, green arrows highlight the clusters where a gene was specifically expressed. The cluster number is indicated for information near to the arrow (see Figure 2A for the annotation of the 21 clusters). The right panels highlight the presence of sNucATAC-seq peaks nearby the TSS of the selected genes. The sc/sNucRNA-seq and sNucATAC-seq clusters are co-annotated to facilitate the comparative analysis. Green boxes and circles highlight major sNucATAC-seq peaks. Regarding At5g65590, a gene not expressed in any root cell type, the red box highlights the lack of open chromatin across the 21 sNucATAC-seq clusters.

**Supplemental Figure 9.**
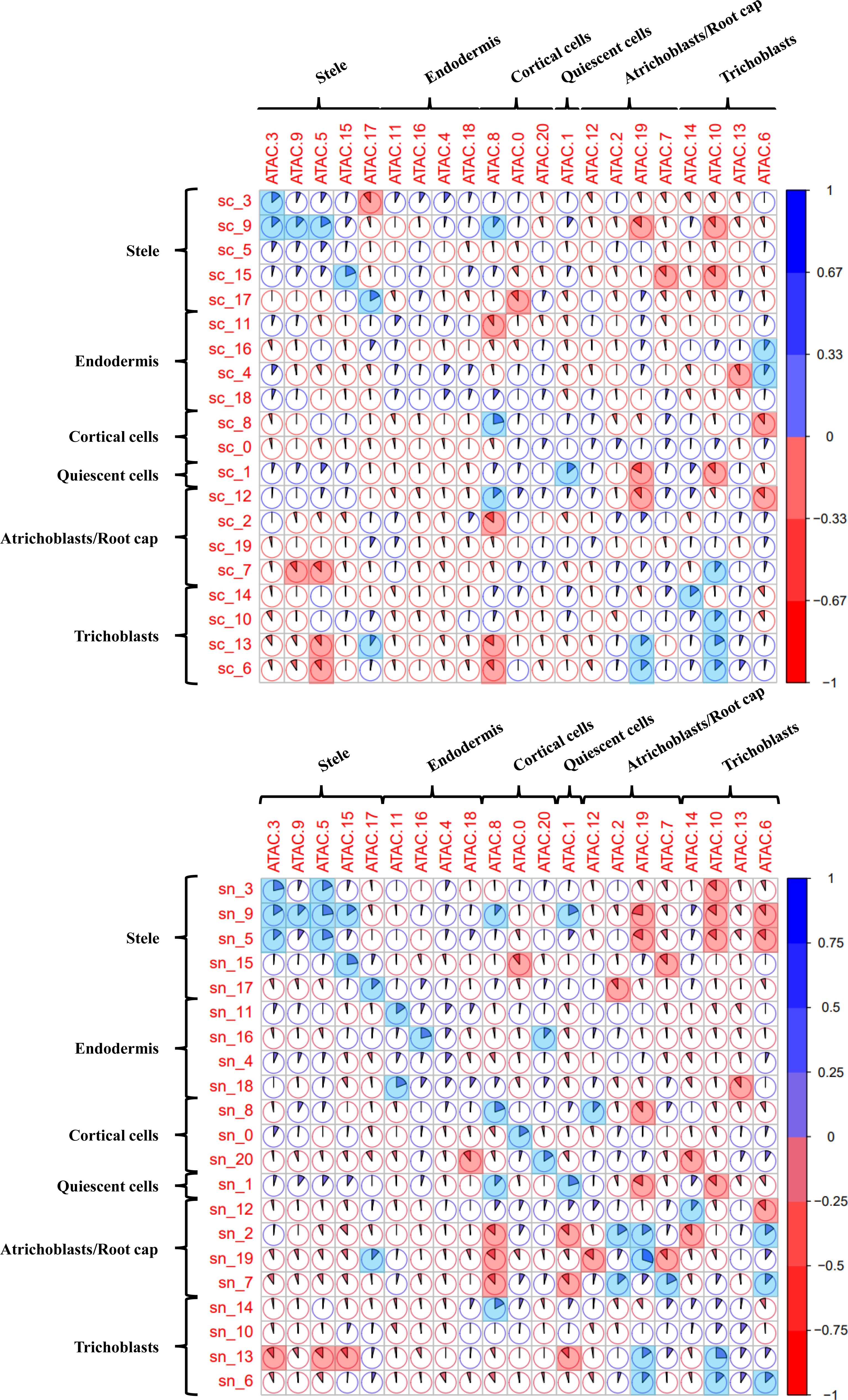
Correlation analyses between gene expression and chromatin accessibility of 628 housekeeping genes for each of the 20 and 21 sc (A) and sNucRNA-seq clusters (B). For each correlation analysis, a Kendall tau-b correlation score was calculated based on the ranking of the cluster according to the expression level of the gene and the level of accessibility of the chromatin fiber (see pies). When significant (p-value < 0.01), positive and negative correlations are highlighted in blue or red, respectively.

**Supplemental Table 1.** Summary of the sequencing of the different single cell and sNucRNA-seq libraries.

**Supplemental Table 2.** List of Arabidopsis root cell-type marker genes.

**Supplemental Table 3.** Expression level and relative accessibility of the chromatin of 337 Arabidopsis root cell-type marker genes.

**Supplemental Table 4.** Expression level and relative accessibility of the chromatin of 628 Arabidopsis root housekeeping genes.

**Supplemental Table 5.** Expression level and relative accessibility of the chromatin of 63 Arabidopsis genes associated with root hair-preferential sNucATAC-seq peaks in their TSS.

